# Aβ-induced synaptic injury is mediated by presynaptic expression of amyloid precursor protein (APP) in hippocampal neurons

**DOI:** 10.1101/2020.07.18.210344

**Authors:** Elena Vicario-Orri, Kensaku Kasuga, Sheue-Houy Tyan, Karen Chiang, Silvia Viana da Silva, Eric A. Bushong, Katherine DeLoach, I-Fang Ling, Liqun Luo, Mark H. Ellisman, Stefan Leutgeb, Edward H. Koo

**Affiliations:** Department of Neurosciences, University of California San Diego, La Jolla, CA 92093, USA; Neurobiology Section and Center for Neural Circuits and Behavior, Division of Biological Sciences, University of California San Diego, La Jolla, CA 92093, USA; National Center for Microscopy and Imaging Research, University of California San Diego, La Jolla, CA 92093, USA; Howard Hughes Medical Institute, Department of Biology, Stanford University, Stanford, CA 94305, USA; Kavli Institute for Brain and Mind, University of California San Diego, La Jolla, CA 92093, USA

**Keywords:** Alzheimer’s disease, LTP, synaptic loss, Aβ

## Abstract

The patterns of Aβ-induced synaptic injury were examined after targeting of the amyloid precursor protein (APP) preferentially to either CA1 or CA3 neurons using Cre-lox technology combined with tetracycline-regulated expression. Both CA1- and CA3-APP-expressing transgenic mouse lines exhibited reduction in long-term potentiation (LTP) only when APP was expressed in neurons presynaptic to the recording site, whereas LTP remained comparable to wild-type mice when APP was expressed in postsynaptic neurons. As quantified by both light and electron microscopy, this orientation-specific impairment in synaptic plasticity was mirrored by synaptic loss in regions receiving axonal inputs from neurons expressing APP. Furthermore, A(plaque deposition also occurred only in the postsynaptic axonal fields of APP-expressing neurons. These deficits were reversed not only with doxycycline to inhibit APP expression but also with γ-secretase and Fyn kinase inhibitors, supporting the interpretation that the observed synaptic injury was mediated by Aβ. Taken together, these results demonstrate that APP/Aβ-induced synaptic toxicity is preferentially initiated by signaling of presynaptically expressed APP to the postsynaptic compartment.

## INTRODUCTION

Alzheimer’s disease (AD), the most common age-related neurodegenerative disorder, is characterized by two pathological hallmarks: intracellular neurofibrillary tangles and extracellular deposits of amyloid-β protein (Aβ). Although these protein aggregates are present in lesser amounts in aged non-demented individuals, the gradual accumulation of brain amyloid is thought to be an early event in the pathogenesis of AD. This viewpoint forms the central tenet of the “amyloid cascade hypothesis”, which states that synapse loss, neuronal dysfunction, neurofibrillary degeneration, and the full manifestations of Alzheimer pathology are initiated by Aβ accumulation in brain [53]. Aβ is derived by sequential cleavage of the amyloid precursor protein (APP), released primarily from neurons in an activity-dependent manner, and subsequently deposited as extracellular aggregates in senile plaques. Recent studies have led to a revision of this hypothesis to emphasize the importance of pre-plaque or soluble oligomeric Aβ species. Specifically, the aggregation state of Aβ peptides is strongly correlated with neurotoxicity: oligomeric but not monomeric Aβ is toxic *in vitro* and *in vivo.* Taken together, many investigators now attribute synaptic toxicity primarily to neuronally released soluble Aβ oligomers [43, 52].

In addition, it has been postulated that Aβ-induced toxicity begins at synapses and that this injury is a major contributor to the cognitive impairments seen in individuals with AD, even before Aβ is deposited extracellularly in plaques [51]. For example, the extent of synapse loss correlates well with the clinical progression of AD [56], as well as severity of dementia, and in the early stages of AD, synapse loss is seen before the development of significant pathology or neuronal death [38, 56, 65]. In line with this concept, APP transgenic mouse lines that develop age-associated amyloid pathology show early reduction in synapses, even prior to the deposition of extracellular Aβ [29, 37, 44]. Addition of Aβ to cultured neurons, cultured hippocampal slices, or injected into the brain results in rapid alterations in synaptic plasticity [46, 64]. Importantly, the reduction in synaptophysin immunoreactivity, electrophysiological abnormalities, and behavioral changes detected before amyloid deposition correlate with increased Aβ levels in brain and may be restored by lowering Aβ levels, suggesting that Aβ-induced synaptic dysfunction is reversible at this stage, at least as modeled in transgenic mice [1, 9, 11, 14, 34].

Although the evidence that Aβ impairs synaptic function in multiple model systems is compelling, there are many crucial details that are unknown. Importantly, the molecular mechanisms by which Aβ causes synaptic injury remain to be clearly elucidated. Many plausible mechanisms have been proposed to explain Aβ toxicity but no clear consensus has emerged. In particular, it is unclear whether Aβ causes synaptic toxicity after release from axonal or dendritic compartments, in no small part owing to the difficulty of addressing this question in the *in vivo* setting. Studies using either cultured primary neurons or organotypic slice cultures have presented evidence supporting the release of Aβ from both axons and dendrites [57, 66]. The Mucke laboratory reported that expression of APP primarily in entorhinal cortex using the tetracycline-regulatable system led to trans-synaptic alterations in the dentate gyrus and CA1 neurons [12]. In this model, amyloid deposits in terminal fields of the perforant pathway originating from the entorhinal cortex suggested that amyloid-associated pathology progresses by presynaptic release. However, the presence of plaques within entorhinal cortex itself indicated that the initial effects could be produced locally by entorhinal cortex neurons. These important findings are also consistent with the spread of neurofibrillary pathology and α-synuclein deposits through the brain, presumably through trans-synaptic connections, as reported by a number of laboratories [16, 22, 35, 63]. Absent from these studies, however, is a clear isolation of the initial toxicity to only afferent or efferent projections such that vulnerability to synaptic injury can be exclusively traced to the pre- or postsynaptic release of Aβ.

In the hippocampus, projections to granule cells of the dentate gyrus (DG) originate from the superficial entorhinal cortex; the DG granule cells then project to CA3 neurons, which then project onto CA1 neurons, which send returning projections to the deep layers of the entorhinal cortex, either directly or indirectly through the subiculum [72]. To address the susceptibility of synapses to Aβ-induced dysfunction from the pre- vs. the post-synaptic compartment, two new lines of transgenic mice were engineered. These two mouse lines use three different transgenes to target APP to either hippocampal CA1 or CA3 neurons by combining the Cre-lox and tetracycline regulated (“tet-off”) systems. This experimental system therefore provides the unique opportunity to electrically stimulate axonal fibers both projecting into and out of CA1 or CA3 neurons, and to record from the dendritic fields of these axonal projections to assess synaptic function. Defects in synaptic plasticity as measured by long term potentiation (LTP) were seen when APP was targeted to presynaptic neurons but not when APP was expressed in postsynaptic neurons, and this pattern was mirrored by synaptic loss and plaque deposition occurring exclusively in compartments postsynaptic to the APP-expressing neurons. The LTP deficits and synaptic loss could be reversed by transgene suppression via doxycycline and γ– secretase inhibitor (GSI) treatment, strongly suggesting a causative role for Aβ-induced synaptic toxicity. Thus, we have leveraged the targeting of APP to two distinct neuronal populations within the hippocampus of transgenic mice to provide evidence that Aβ released from pre-synaptic compartments preferentially results in synaptic dysfunction.

## RESULTS

### Characterization of Site-specific APP Expression in CA1 or CA3 Triple Transgenic Mice

In this study, we wished to direct APP expression preferentially to either neurons in the CA1 or CA3 region and assess perturbations in synaptic function. To achieve this spatial and temporal control of APP expression *in vivo*, we combined the Cre/loxP and tet-off systems to generate two lines of mice (hereon designated as “CA1-APP’ or “CA3-APP” mice), each carrying three transgenes to enable this targeting to CA1 or CA3 neurons, respectively. The three separate transgenic lines used in our breeding scheme are: 1) CAG promoter driving tetracycline transactivator (tTA) situated downstream of a lox-stop-lox cassette (“Tg1:ZtTA” in Fig. 1A) [33], 2) CA1- or CA3-Cre transgenic mouse lines (“Tg2” in Fig. 1A) [39, 58], and 3) TRE-APP where the human APP transgene (“Tg3” in Fig. 1A), encoding the “Swedish” and “Indiana” mutations, is controlled by the tetracycline responsive promoter element (TRE) [21]. In this tet-off system, the presence of doxycycline (dox) is able to silence APP expression. In a mouse carrying all three transgenes, APP will be expressed in a cell-specific manner upon activation by tTA, which itself can only be expressed upon Cre-mediated excision. To direct hippocampal-specific expression, either the α-CaMKII (T29-1 line) or Grik4 promoter directed-Cre expression in CA1 or CA3 neurons, respectively [39, 58].

**Fig. 1.**
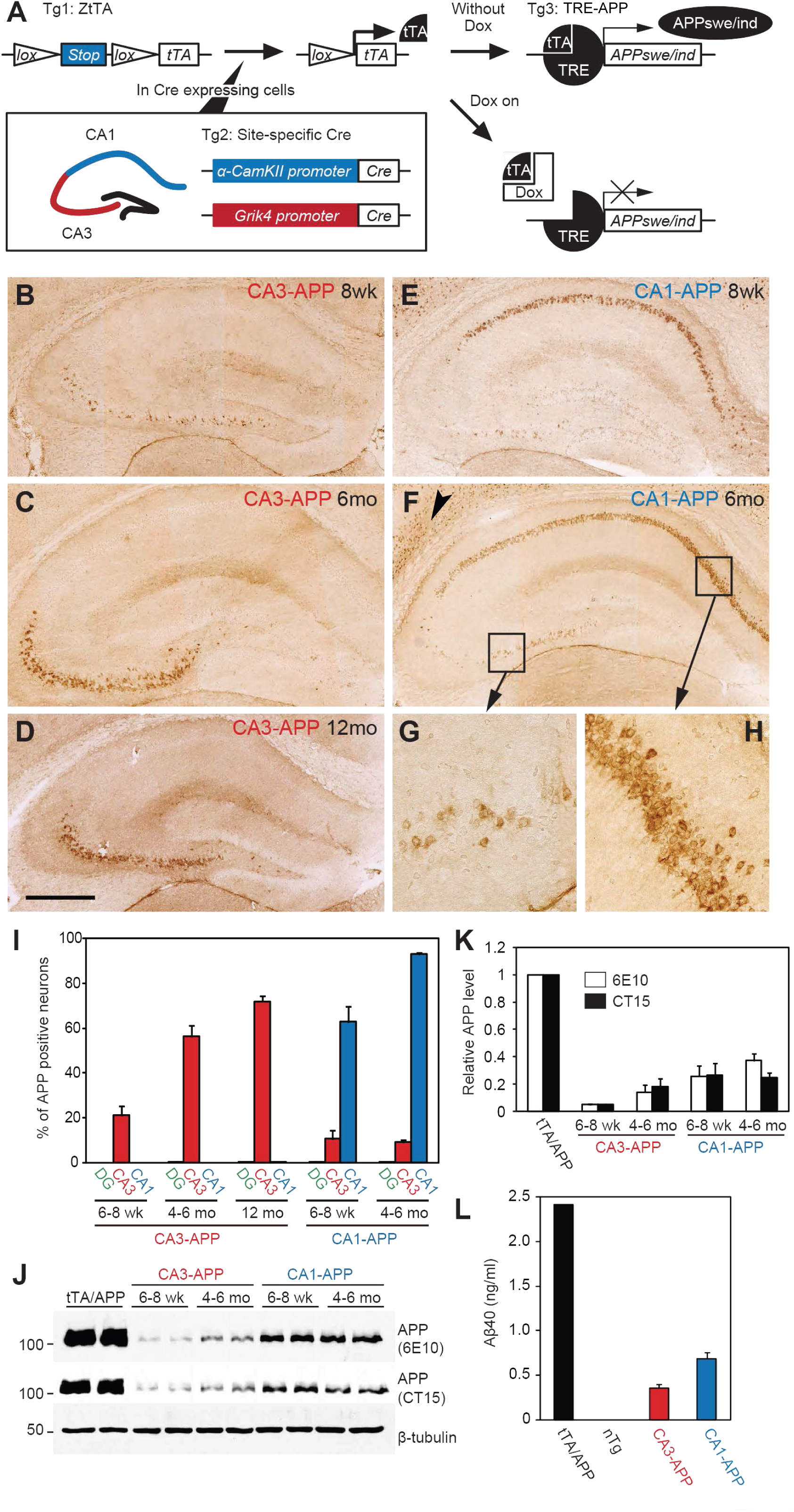
Site-specific Expression of Transgenic APP in CA3- and CA1-APP Mice. **(a)** Schematic of breeding scheme. Transgene 1 (Tg1): ZtTA, tetracycline-controlled transactivator protein (tTA) transgenic mouse line under the control of Cre-loxP recombination. Tg2: Site-specific Cre transgenic mouse lines under the regulation of α-CaMKII promoter (T29-1) for CA1 (blue) and Grik4 promoter for CA3 (red) expression, respectively. Tg3: TRE-APP, an APP transgenic mouse line under the control of a tetracycline-responsive promoter element (TRE). When all three transgenes are present, APP expression is limited to either CA1 or CA3 neurons as specified by the Cre promoter (Tg2), and can be suppressed by doxycycline administration (Dox). **(b-d)** Representative photomicrographs of coronal sections from CA3-APP mice at 8 weeks (b), 6 months (c), and 12 months of age (d), stained for APP expression with anti-human Aβ antibody 6E10. Note specificity of APP expression in CA3 neurons only (brown). **(e-f)** Representative photomicrographs of coronal sections from CA1-APP mice at 8 weeks (e) and 6 months (f) stained with 6E10. The arrowhead indicates APP expression in a region of the cortex; higher magnification images of CA3 (g) and CA1 regions (h) from panel (f) indicate sparse immunostaining of APP in a subset of CA3 neurons compared to much more robust, extensive expression in CA1 neurons. Scale bar in B-F: 0.5 mm. **(i)** Quantification of percent APP-positive neurons within the DG (green bars), CA3 (red bars), and CA1 (blue bars) regions in CA3- and CA1-APP mice from 6-8 weeks up to 12 months of age. The percentage of APP immunopositive neurons increased in an age-related manner in CA3-APP mice, but APP expression remained absent in DG or CA1 regions. In CA1-APP mice, APP immunopositivity increased from ∼60% to 90% of CA1 with age, but remained at ∼10% in CA3 neurons. Data are averages of 3 to 4 mice from each age group (±SEM). **(j)** Representative APP immunoblots from dissected hippocampi of mice from two age groups. **(k)** Quantification of the immunoblots showed that levels of APP were approximately 5% (6-8 week-old CA3-APP), 15% (4-6 month-old CA3-APP), 25% (6-8 week-old CA1-APP), and 35% (4-6 month-old CA1-APP) of the amount found in 2 month-old α-CaMKII-tTA×TRE-APP (tTA/APP) mice. Data were averages of three independent experiments (±SEM). **(l)** ELISA-based quantification of Aβ40 levels from SDS-extracted hippocampal lysates of two month-old transgenic mice (n=3 ± SEM).

The presence of APP expression in the expected neuronal populations in adult mice was confirmed by immunohistochemistry using a human-specific APP (6E10) antibody (Fig. 1B-F). In CA3-APP mice, APP expression was essentially restricted to CA3 neurons in the hippocampus. Further, this expression increased in an age-dependent manner, increasing from approximately 20% to 80% of CA3 neurons from 8 weeks to 12 months of age (Figures 1B-D and 1I). Rare scattered neurons in the neocortex and amygdala were also immunopositive for APP (data not shown). We also confirmed that APP expression was not present in adjacent CA2 neurons, which are morphologically very similar to CA3 cells, by double labeling with human APP and the CA2 marker Purkinje cell protein 4 (PCP4) [27, 32]. The staining indicated that transgenic APP expression in CA3-APP mice was absent from CA2 neurons and confined to CA3 neurons (Supplementary Fig. 3A). In CA1-APP mice, APP was preferentially expressed in CA1 neurons, again in an age-dependent manner: increasing from expression in approximately 60% to over 90% of CA1 neurons from 8 weeks to 6 months of age (Fig. 1E, F, H and I). A minor percentage of CA3 neurons (<10%) were also immunopositive for APP and remained at this low level up to 6 months of age (Fig. 1G, I). From around 6 months onwards, APP expression was diffusely present in other cortical regions (arrowhead in Fig. 1F), matching previously reported expression patterns for the commonly used forebrain α-CaMKII promoter driven tTA mice [10, 19, 71]. Spatial specificity was thus not maintained in CA1-APP mice as they aged and all analyses were therefore limited to mice under 6-7 months of age. However, APP immunostaining was not detected in either dentate granule cells or in glia (data not shown).

By Western blotting, expression of transgenic APP was readily detected in dissected hippocampi and increased in an age-dependent manner (Fig. 1J, K). Further, APP levels were approximately two-fold higher in CA1-APP than CA3-APP mice at 4-6 months of age (Fig. 1J, K), consistent with the more numerous APP immunopositive neurons in the CA1-APP mice (Fig. 1I). Soluble Aβ40 levels from SDS-extracted hippocampal lysates measured by the MSD assay system at 8 weeks of age were also approximately two-fold higher in CA1-APP mice as compared to CA3-APP mice (Fig. 1L). As expected, levels of both APP and Aβ40 were substantially lower in both mouse lines as compared to non-selective forebrain- and hippocampus-directed APP expression in the α-CaMKII-tTA×TRE-APP mice analyzed in parallel as a positive control [21]. These results demonstrated excellent localization of APP expression to the two desired neuronal populations.

### Pre-synaptic APP Expression Caused LTP Deficits

Having established the predicted spatial localization of APP transgene expression to CA1 or CA3 neurons in the two APP transgenic mouse lines, we then asked if synaptic plasticity is impaired when APP is expressed in the pre- or postsynaptic neurons. Specifically, one attractive property of basing our current experimental system in the hippocampus, which can mostly be considered as a monosynaptic pathway, is the opportunity to assess the synaptic activity of both efferent and afferent projections of the same neuronal populations. We first considered the hypothesis that Aβ release from presynaptic terminals alters postsynaptic function, and accordingly predicted that stimulation of axonal fibers from APP-expressing CA3 or CA1 neurons would demonstrate changes in synaptic measurements for synapses onto CA1 or subicular dendrites, respectively. First, LTP, basal synaptic transmission, and paired-pulse facilitation (PPF) were assessed in CA3-APP mice following stimulation of Schaffer collaterals and recording from the CA1 dendritic field, hereon designated as the SC-CA1 synapse (Fig. 2A). In this paradigm of presynaptic APP expression, SC-CA1 LTP was reduced at both 6-8 weeks (Supplementary Fig. 1A) and 4-6 months (Fig. 2B) when compared to age-matched control littermates. However, at this age there were no changes in either basal synaptic transmission (Fig. 2C) or PPF tested with interpulse intervals from 50-400 ms (Supplementary Fig. 1B), the latter reflecting unaltered pre-synaptic vesicle release probability. Given the lack of changes in PPF regardless of interpulse intervals, tests of PPF in later experiments used only the 50 ms interpulse interval. Basal synaptic transmission, as measured by fEPSP input-output (I/O) curves, was found to be depressed only in older 12 month-old CA3-APP mice (Fig. 2D), while PPF remained unchanged as compared to controls at this age (Supplementary Fig. 1D). Second, we also performed synaptic measurements in 6-8 week-old (data not shown) and 4-6 month-old CA1-APP mice, now stimulating the CA1 axons and recording in the subiculum, hereon designated as the CA1-SUB synapse (Fig. 2E). Remarkably, essentially the same results were seen in this CA1-APP mouse line, namely, significant reduction in CA1-SUB LTP (Fig. 2F) but preserved basal synaptic transmission (Fig. 2G) and PPF (Supplementary Fig. 1E). The results from the CA3-APP mice and the CA1 mice suggested that APP/Aβ originating from presynaptic neurons and released from presynaptic sites of two different neuronal populations impaired LTP without altering the pre-synaptic transmitter release probability or basal synaptic transmission up to 6 months of age in both mouse lines.

**Fig. 2.**
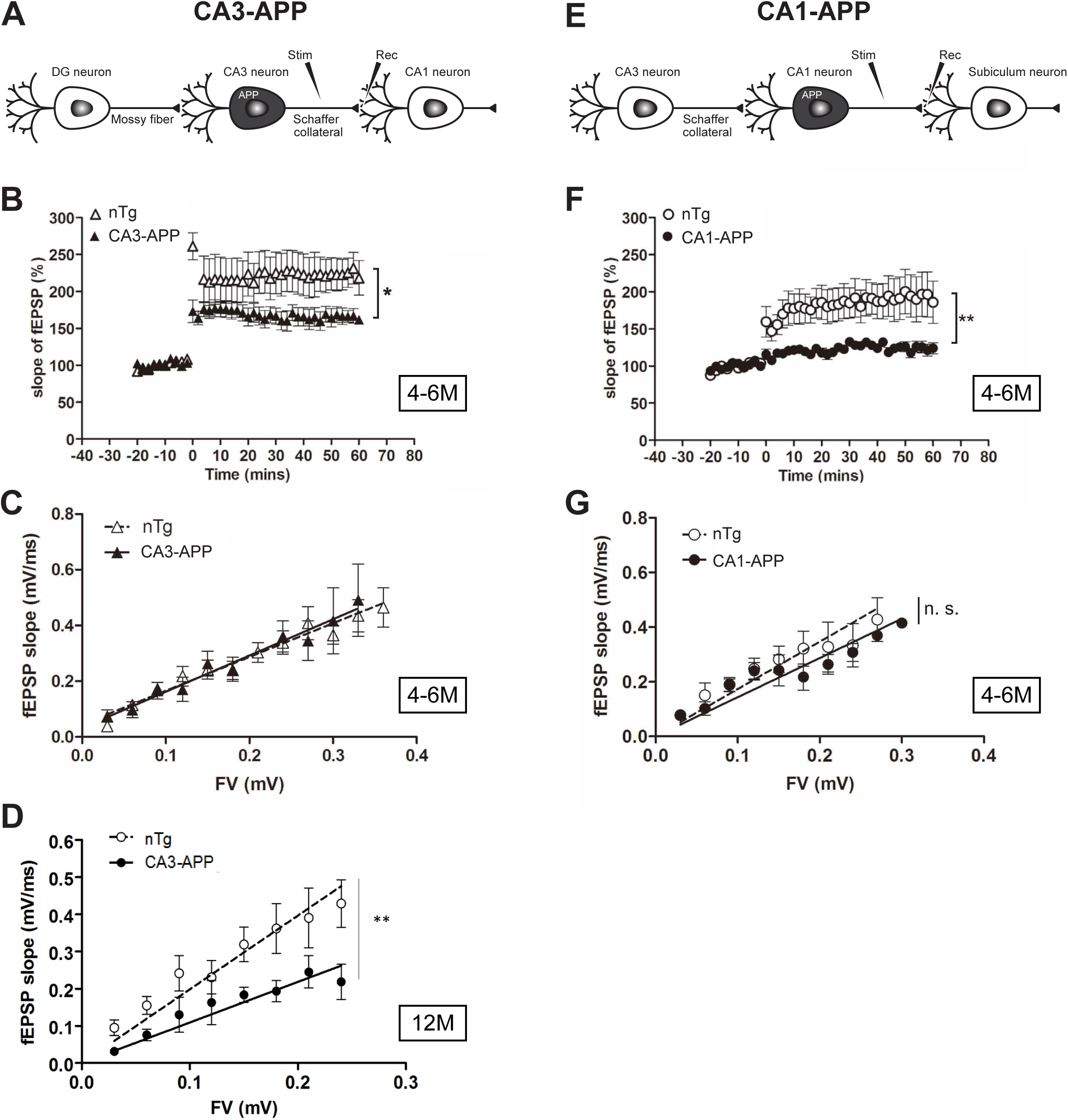
Pre-synaptic APP Expression Impaired Synaptic Plasticity in CA3- and CA1-APP Mice. **(a)** In CA3-APP mice, field excitatory postsynaptic potentials (fEPSPs) were measured in CA1 stratum radiatum following stimulation of Schaffer collaterals (SC-CA1 LTP). At the recording site, the stimulated axons (Schaffer collaterals) from APP-expressing CA3 cells (black) are presynaptic, and CA1 neurons without APP expression are postsynaptic. **(b)** CA3-APP mice (black triangles) showed significant SC-CA1 LTP reduction compared to control non-transgenic littermates (nTg, open triangles) at 4-6 months of age (ANOVA and *post hoc* Tukey’s test, *p<0.05; nTg: n=10 slices from 5 mice; CA3-APP: n=13 from 7 mice). **(c)** At 4-6 months of age, there were no significant changes in basal synaptic transmission, as indicated by a matching slope between the fiber volley amplitude (FV) and the slope of field EPSP (fEPSP) (p>0.5; nTg: n=15 slices from 6 mice; CA3-APP: n=15 from 6 mice). **(d)** Basal synaptic transmission was impaired in 12 month-old CA3-APP mice, as indicated by a lower fEPSP slope in transgenic mice for each corresponding fiber volley (FV) amplitude value. Slope for CA3-APP: 1.06±0.5, slope for nTg: 1.97±0.10 (** p<0.01; n=7 mice/group). **(e)** In CA1-APP mice, APP was expressed in presynaptic CA1 neurons and field potentials were recorded in the substratum radiatum of the subiculum following stimulation of CA1 axons projecting to subiculum (CA1-SUB LTP). **(f)** CA1-APP mice (black circles) showed CA1-SUB LTP impairment as compared to control nTg littermates (open circles) at 4-6 months of age. (**p<0.01; nTg: n=12 slices from 7 mice; CA1-APP: n=17 from 7 mice). **(g)** Basal synaptic transmission was normal at the CA1 to subiculum synapses in 4-6 month-old CA1-APP mice (p=0.38; nTg: n=13 slices from 7 mice; CA1-APP: n=14 from 7 mice).

### Post-synaptic APP Expression Did Not Reduce LTP

We next asked whether synaptic function is altered when APP was expressed in the postsynaptic neuron as opposed to the presynaptic neuron, as analyzed previously. In CA3-APP mice, mossy fibers (MF) projecting from dentate gyrus were stimulated and recordings taken at the MF to CA3 synapses, hereon designated as MF-CA3 (Fig. 3A) and in CA1-APP mice, at the SC-CA1 synapses (Fig. 3F). In marked contrast to presynaptic APP expression, MF-CA3 LTP in CA3-APP mice was unchanged as compared to non-transgenic littermate controls at 4-6 months of age (Fig. 3B) or 12 months of age (Fig. 3D). SC-CA1 LTP was also not perturbed in CA1-APP mice as compared to littermate controls at 6-8 weeks (data not shown) and at 4-6 months of age (Fig. 3G). Basal synaptic transmission (Fig. 3C, E, & H) and PPF (Supplementary Fig. 2) were not altered in either CA3- or CA1-APP mice at the respective synapses. Taken together, these results from two different neuronal populations demonstrated that synaptic plasticity was impaired only when APP was expressed in presynaptic but not postsynaptic neurons.

**Fig. 3.**
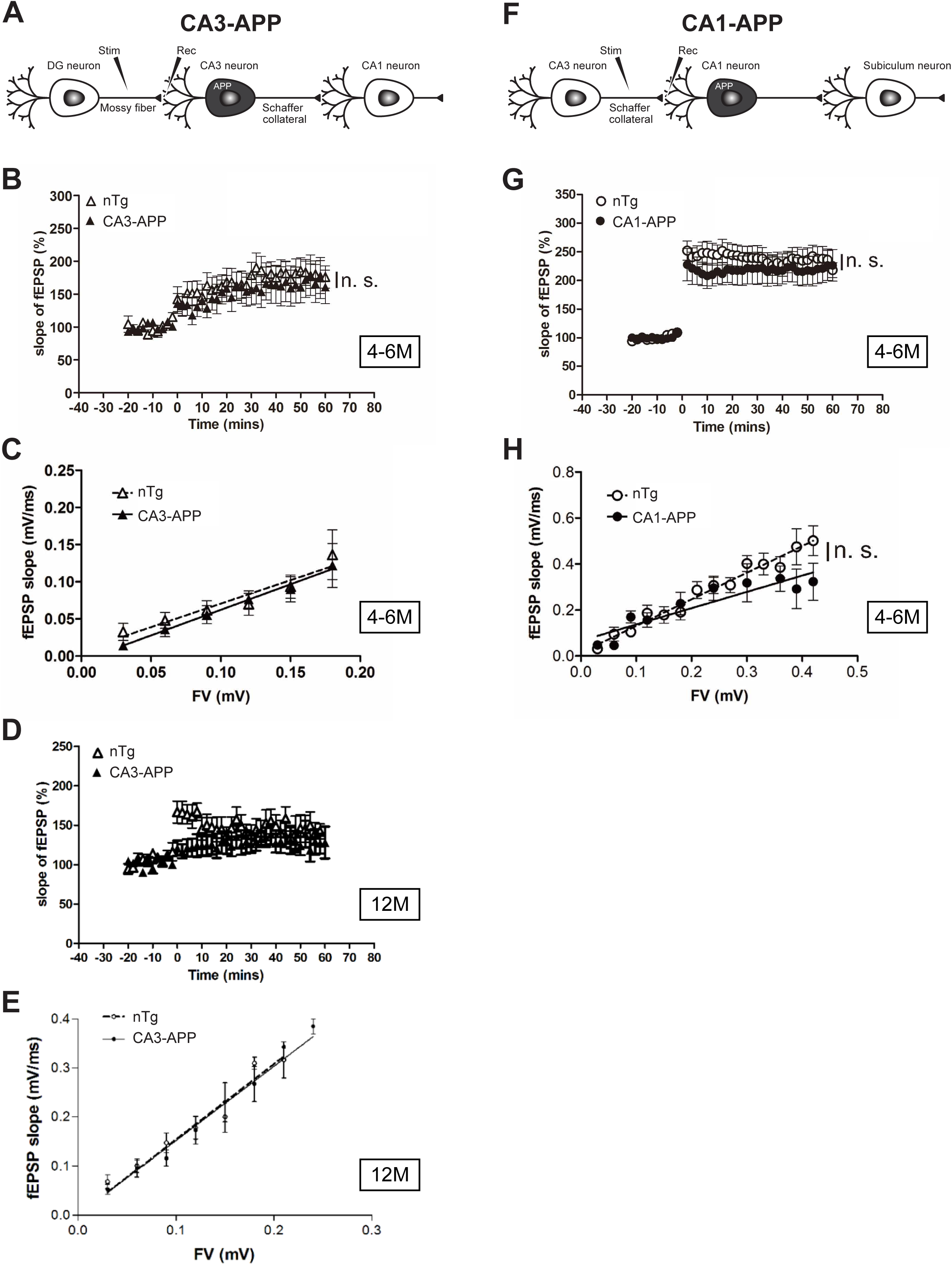
Post-synaptic APP Expression Did Not Impair Synaptic Plasticity. **(a)** APP was expressed in CA3 neurons (black) and field potentials were recorded from the CA3 region, which is postsynaptic following stimulation of mossy fiber projections from dentate gyrus (MF-CA3 LTP). **(b)** CA3-APP mice (black triangles) showed normal MF-CA3 LTP as compared to control nTg littermates (open triangles) at 4-6 months of age (ANOVA and *post hoc* Tukey’s test, p>0.5, nTg: n=10 slices from 5 mice; CA3-APP: n=11 from 6 mice). **(c)** No significant changes in basal synaptic transmission were observed (p>0.5; nTg: n=10 slices from 6 mice; CA3-APP: n=9 from 5 mice). **(d)** 12 month-old CA3-APP mice (black triangles) showed normal MF-CA3 LTP as compared to controls (open triangles). **(e)** No significant changes in basal synaptic transmission were observed in 12 month-old mice. **(f)** APP was expressed in CA1 neurons (black) and field potentials were recorded from CA1 stratum radiatum, which is postsynaptic following stimulation of Schaffer collaterals (SC-CA1 LTP). **(g)** CA1-APP mice (black circles) demonstrated normal SC-CA1 LTP as compared to control non-transgenic littermates (open circles) at 4-6 months of age (p>0.5; nTg: n=12 slices from 6 mice; CA1-APP: n=11 from 5 mice). **(h)** No significant changes in basal synaptic transmission were observed (p>0.5; nTg: n=16 slices from 7 mice; CA1-APP: n=11 from 5 mice).

### Pre-synaptic and Not Post-synaptic APP Expression Caused Synaptic Loss

Given the known synaptotoxic effects of Aβ [26], we next looked for evidence of synapse loss in CA3- and CA1-APP mice. In light of the reduction in LTP we observed when APP was expressed presynaptically, we hypothesized that the reduction in synapse density would predominantly occur in the dendritic fields receiving axonal projections from APP expressing neurons. The axon terminals of hippocampal cells are well-defined: in brief, the mossy fiber projections of DG granule cells terminate in the CA3 stratum lucidum (CA3 SL, orange; Fig. 4A), the Schaffer collaterals of the CA3 neurons project to the CA1 stratum radiatum and stratum oriens (CA1 SR and CA1 SO, blue), while the CA1 axonal projections terminate in the subiculum. CA3 neurons also send recurrent projections to other CA3 neurons in the CA3 stratum radiatum (CA3 SR, blue). To quantify synaptic numbers, we used the percent surface area of synaptophysin and PSD-95 immunoreactivity as a measure of pre- and post-synaptic compartments, respectively [3, 5, 20, 67]. Loss of synaptophysin- and PSD95-immunoreactive puncta has been reported in other APP transgenic mouse lines and is a characteristic feature of AD pathology [2, 37, 55, 56]. At 6 months of age in CA3-APP mice, there were no detectable changes in synaptic density as assessed by synaptophysin or PSD-95 immunostaining in the CA1 SR and CA1 SO layers where CA3 neurons send their axonal projections (Fig. 4B, Supplementary Fig. 3B), even though mice at this age already exhibited LTP deficits (Fig. 2B).

**Fig. 4.**
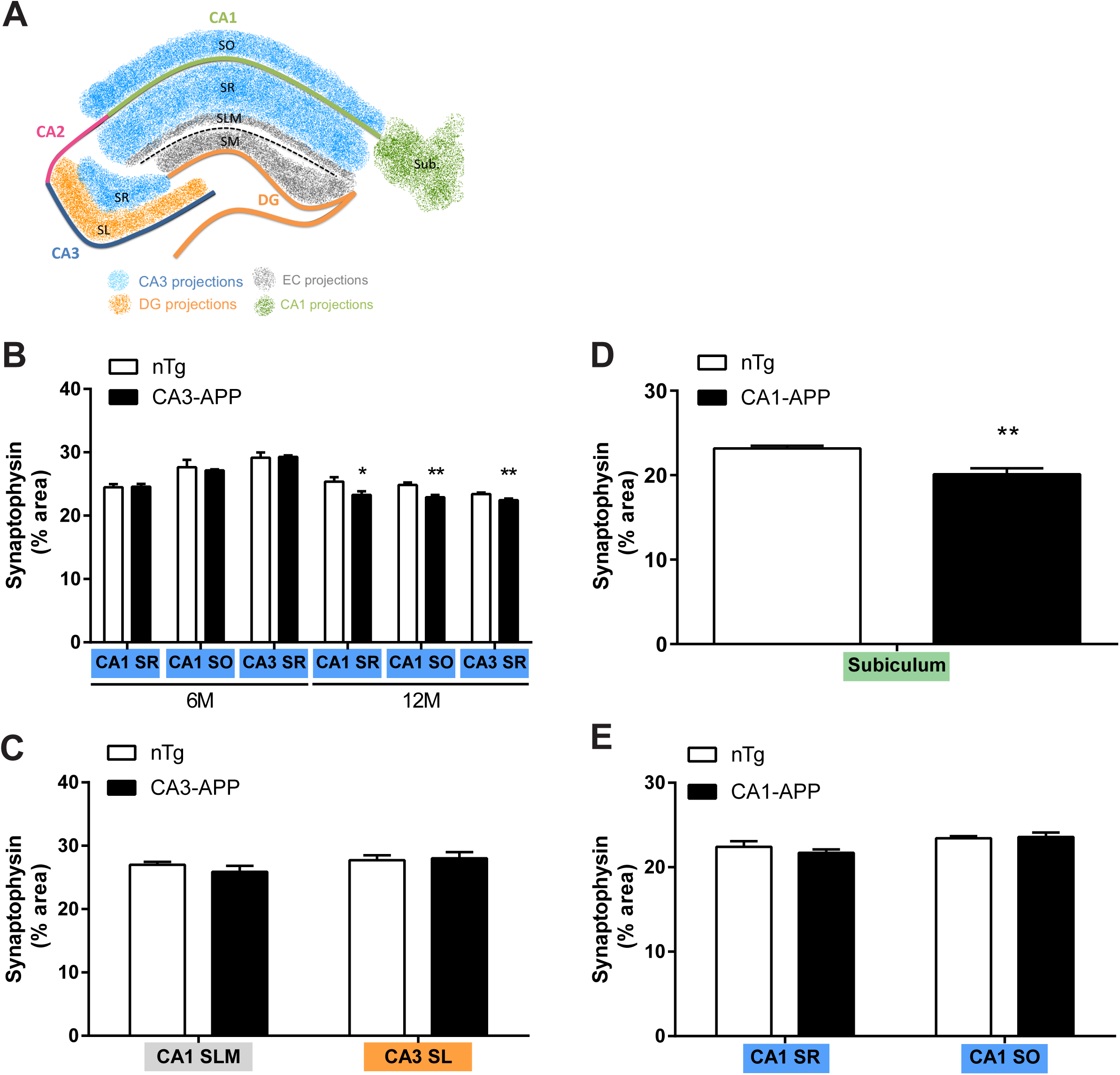
Synaptic loss in CA3- and CA1-APP mice only occurs in regions receiving axonal projections from APP-expressing neurons. **(a)** Diagram of termination zones of hippocampal projections. Granule cells of the dentate gyrus (DG) project to CA3 and their efferent axons, also known as mossy fibers, terminate in an area called the stratum lucidum (CA3 SL, orange). In turn, CA3 pyramidal neurons project, via Schaffer collaterals, to the stratum radiatum (CA1 SR, blue) and to the stratum oriens (CA1 SO, blue), where they synapse onto dendrites of CA1 neurons. CA3 neurons also send recurrent projections to other CA3 cells, with synapses in the CA3 stratum radiatum (CA3 SR, blue). CA1 also receives axonal projections via the perforant path from neurons in the enthorhinal cortex (EC), and these terminals synapse in the stratum lacunosum-moleculare layer (CA1 SLM, gray). Finally, CA1 neurons send projections to subiculum (Sub., green). **(b)** Percent area of synaptophysin immunoreactivity in CA1 SR, CA1 SO and CA3 SR in 6 and 12 month-old CA3-APP mice. 12 month-old CA3-APP mice had decreased immunoreactivity in all three termination zones of CA3 projections compared to control littermates. (All values are mean ± SEM; n=9 mice/group. Unpaired two-tailed t-test, *p<0.05, ** p<0.01, ***p<0.001; CA1 SR: p=0.03, CA1 SO: p=0.002, CA3 SR: p=0.008.) **(c)** Percent area of synaptophysin immunoreactivity in CA1 SLM and CA3 SL of 12 month-old CA3-APP mice was unchanged compared to control littermates. **(d)** 6 month-old CA1-APP mice exhibited a significant decrease of synaptophysin immunoreactivity in the subiculum, where CA1 neurons send axonal projections, compared to nTg. (n=8 mice/group, p=0.004.) **(e)** In 6 month-old CA1-APP mice, synaptophysin immunoreactivities in the areas where APP-expressing CA1 neurons do not send axonal projections were unchanged compared to nTg mice.

However, in 12 month-old CA3-APP mice, small (∼10%) but significant decreases in both synaptophysin and PSD-95 immunostained puncta were observed (Fig. 4B, Supplementary Fig. 3B), notably at an age where we first observed depression of basal synaptic transmission (Fig. 2D). Furthermore, we also noted a minor but significant reduction in the CA3SR layer (Fig. 4B and Supplementary Fig. 3B), wherein CA3 recurrent collateral axons synapse back on dendrites of CA3 neurons. Note however that the reduction seen by synaptophysin and PSD-95, although statistically significant, was quite small (∼5%, unpaired, two-tailed t-test), approximately half the reduction that was observed in the CA1 dendritic fields.

The distinct innervation patterns into the hippocampus are such that dendritic fields that receive inputs from neurons not expressing APP would not be expected to exhibit synaptic loss, thus serving as an important negative control. Directly adjacent the CA1 SR area lies the CA1 stratum lacunosum-moleculare (CA1 SLM) layer, which receives projections primarily from layer III neurons of the entorhinal cortex via the perforant pathway (grey areas in Fig. 4A), but not from CA3 neurons. Indeed, as hypothesized, synaptophysin and PSD-95 immunoreactivity was unchanged in CA1 SLM of CA3-APP mice compared to control littermates (Fig. 4C and Supplementary Fig. 3C). Furthermore, the CA3 stratum lucidum (CA3 SL), clearly demarcated from the CA3 SR by the presence of the mossy fiber giant boutons, also did not exhibit any decrease in synaptophysin or PSD-95 staining (Fig. 4C, Supplementary Fig.3C), as their inputs originate from the dentate gyrus.

Consistent with what we found in the 12 month-old CA3-APP mice, synaptophysin and PSD-95 immunoreactivity in 6 month-old CA1-APP mice was comparably decreased in the subiculum, which receives axonal projections from CA1 neurons (Fig. 4D). Further, the CA1 SR and CA1 SO regions, which receive projections from CA3 neurons not expressing APP, showed no indication of synaptic loss (Fig. 4E). Thus, the regions of synapse loss as measured by synaptophysin and PSD-95 densities were in the axonal fields of neurons expressing APP in both CA1- and CA3-APP mice.

To further confirm that the loss of immunostained synaptic markers accurately reflects actual loss of synapses, we performed synaptic counts on serial block-face scanning electron microscopy (SBEM) images taken from 12 month-old CA3-APP mice (Fig. 5A). In line with the previous results using immunostaining of synaptic markers, synaptic counts as assessed by SBEM in the CA1 SR, CA1 SO, and CA3 SR layers were significantly reduced as compared to non-transgenic littermates (Fig. 5B). Further, the magnitude of reduction was two-fold larger than that quantified from immunostaining of synaptic markers. As predicted, the synaptic counts in the CA1 SLM (EC projections) and CA3 SL (DG projections) regions showed no differences (Fig. 5C), again consistent with the synaptophysin and PSD-95 immunoreactive densities. Taken together, the results from light and electron microscopy in CA3- and CA1-APP mice argue strongly that synaptic damage only occurs in dendritic layers that receive projections from APP-expressing cells.

**Fig. 5.**
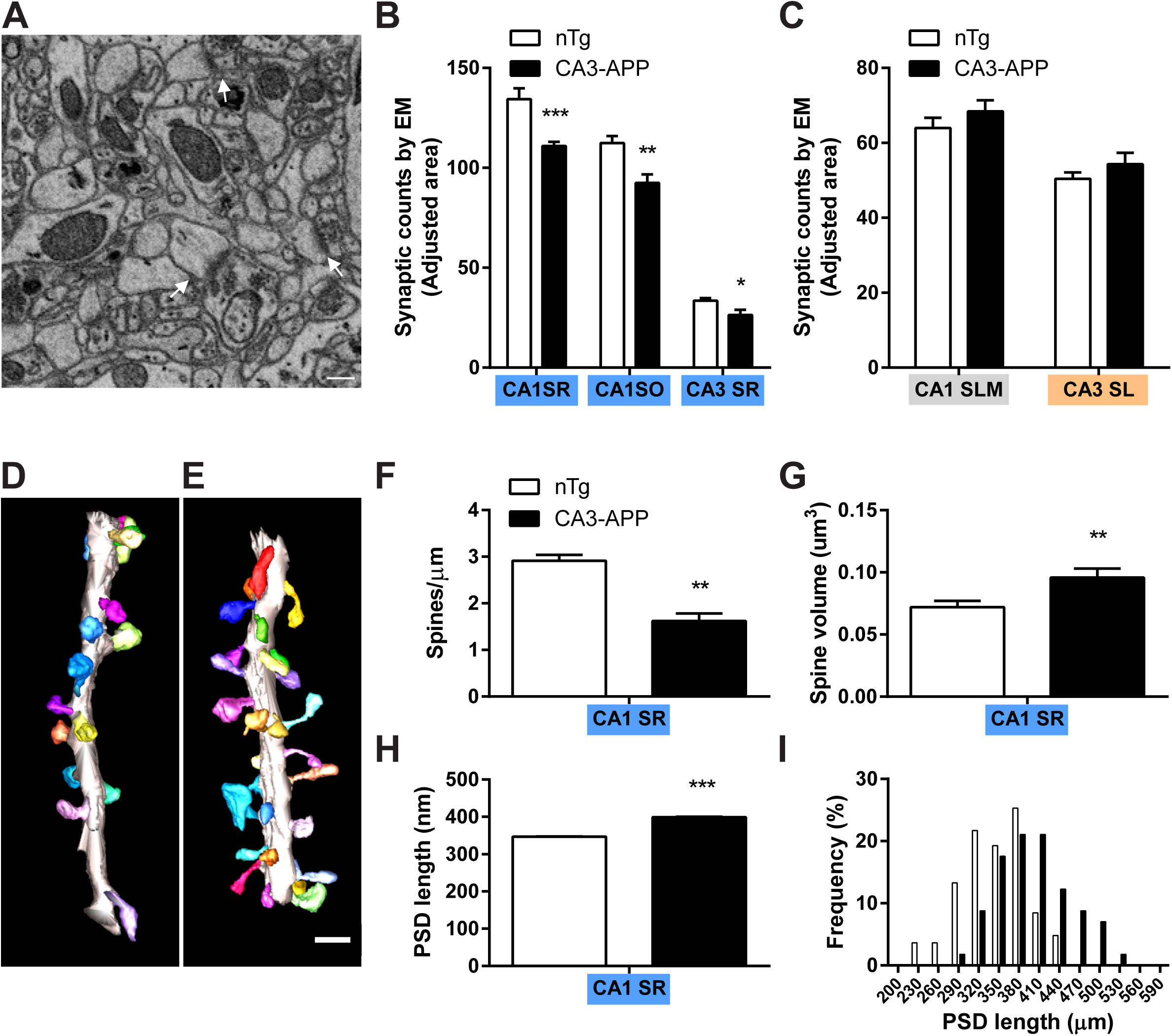
Electron microscopy confirms synaptic loss and decreased synaptic density in axonal terminal fields of APP-expressing neurons in CA3-APP mice. **(a)** Representative SBEM image of CA1 SR in a 12 month-old CA3-APP mouse. Arrows indicate examples of synapses. Scale bar: 0.5 µm. **(b)** Synapse counts of SBEM images per adjusted area. 12 month-old CA3-APP mice have a significantly decreased number of CA1 SR, CA1 SO, and CA3 SR synapses compared to nTg mice. (All values are mean ± SEM; n=80-100 spines from 3-4 dendrites, 2 mice/group. Unpaired two-tailed t-test, *p<0.05, *p<0.01, ***p<0.001; CA1 SR: p=0.0005, CA1 SO: p=0.0017, CA3 SR: p=0.022.) **(c)** 12 month-old CA3-APP mice have similar synapse counts in the CA1 SLM and CA3 SL as compared to control littermates. **(d-e)** Representative reconstruction of dendritic segments (gray) and associated spines (colored) in CA1 SR in a 6 month-old CA3-APP mouse (D) and a nTg littermate control (E). Scale bar: 1 µm. **(f)** Quantification of the number of spines per length of dendrite in the CA1 SR shows that 6 month-old CA3-APP mice have a significantly lower number of spines per µm. (*p<0.05, **p<0.01, ***p<0.001 unpaired two-tailed t-test; n=80-100 spines from 3-4 dendrites, 2 mice/group.) **(g)** Quantification of spine volume in the CA1 SR showed a significant increase in the 6 month-old CA3-APP mice compared to the nTg. (CA3-APP: n=54, nTg: n=89 spines from 2 mice, p=0.007.) **(h)** Length of the PSD in CA1 SR synapses was significantly increased in 6 month-old CA3-APP mice. (CA3-APP: n=57, nTg: n=83 synapses from 2 mice, p<0.0001.) **(i)** Distribution of CA1 SR synaptic PSD lengths in 6 month-old nTg and CA3-APP mice showed a shift in CA3-APP mice towards increased length.

Given that we found no apparent synaptic loss in young (6 months old) CA3-APP mice by synaptophysin and PSD-95 staining despite significant reduction in synaptic plasticity at this age, we wondered whether we could detect morphological changes at the synaptic level prior to frank synaptic loss. Accordingly, SBEM coupled with 3D reconstruction was used to analyze the CA1 dendritic layers in 6 month-old mice, as this method offered higher resolution for studying alterations in synapse structure [68, 69]. The 3D reconstruction was undertaken from SBEM images and four dendritic shafts were examined in the CA1 SR layer where synaptic loss was previously detected in 12 month-old CA3-APP mice (Fig. 5B). Volumetric reconstruction of the dendritic segments and associated spines did not reveal any evidence of overt structural abnormalities, but surprisingly, the density of spines was significantly reduced by almost 50% in the dendrites of CA3-APP compared to control littermate mice (Fig. 5F). This reduction is particularly notable not only for the greater magnitude in spine reduction as compared to the previous quantification by immunofluorescence (Fig. 4B), but also for being detectable in the younger, 6 month-old animals. A further and unexpected finding is an increase (∼30%) in the volume of dendritic spines, suggesting that the remaining spines were, on average, larger than control littermates’ (Fig. 5G). To confirm this observation, the length of the postsynaptic densities (PSDs), another measure of synaptic size and presumably synaptic transmission strength, was assessed. This analysis showed that the length of postsynaptic densities (PSDs) was also significantly increased in CA3-APP mice (Fig. 5H-5I).

### Plaque Accumulation Is Only Present in the Terminal Fields of APP-Expressing Neurons

With a few exceptions, APP transgenic mouse lines to date have generally been driven by neuron-specific promoters, such as Thy-1 or PDGF, or in a more generalized CNS-wide manner, such as with the PrP promoter. In these mice, APP expression is generally diffuse in brain and the resulting Aβ deposits are located in the neocortex and hippocampal areas, much as seen in human brain. However, the cells from which the deposited Aβ originated cannot be precisely determined, in contrast to our newly developed models, in which APP expression is restricted to certain neuronal populations. We therefore examined the presence and location of Aβ deposits in CA3- and CA1-APP mice. In CA3-APP mice, no Aβ deposits were seen in mice up to 9 months of age. In a subset of 12 month-old animals (Table 1), rare and infrequent deposits, always of the diffuse variety, were present as solitary deposits, but only in the terminal fields of CA3 to CA1 projections, i.e., CA1 SR and SO (Fig. 6A and Supplementary Fig. 4E-F). This pattern of plaque accumulation, i.e., in the known targets of CA3 axons, strongly implicated the release and subsequent deposition of Aβ from axonal terminals, as demonstrated recently using an optogenetics approach [75]. Importantly, because there was little spreading of APP expression beyond the region of CA3 terminals in the CA3-APP mice at this age, the origin of Aβ can be attributed to CA3 neurons with some certainty. Interestingly, virtually all Aβ deposits were present in posterior hippocampal regions. Compared to CA3-APP mice, Aβ deposits were detected in CA1-APP mice much earlier and in a clear age-associated manner (Fig. 6B, Table 1). In 6-7 month-old mice, Aβ plaques were detected only in the subiculum but not within the CA1 region itself at a time when APP immunoreactivity had already spread to limbic and neocortical areas (Fig. 6B and Supplementary Fig. 4A-B, Fig. 1I). Because scattered neurons in the subiculum were also positive for APP by immunostaining, the possibility of dendritic release of Aβ from subicular neurons cannot be excluded. Notably however, Aβ deposits were not detected in dendritic fields of CA1 neurons, in spite of the presence of APP expression in most CA1 neurons by immunohistochemistry. In mice older than 6-7 months, amyloid deposits were always prominent in the subiculum as compared to the hippocampus and there were plaques irregularly scattered throughout the cortex concomitant with widespread expression of APP (Supplementary Fig. 4C-D), consistent with the activation of transgene expression in other neuronal populations as reported [10, 19, 71]. In sum, the observations from these two APP transgenic mouse lines with preferential expression of APP in CA1 or CA3 neurons provided strong support that Aβ deposits originated primarily from Aβ released from axon terminals of presynaptic neurons rather than from local or dendritic release, as hypothesized by a number of prior studies [7, 48, 66].

**Fig. 6.**
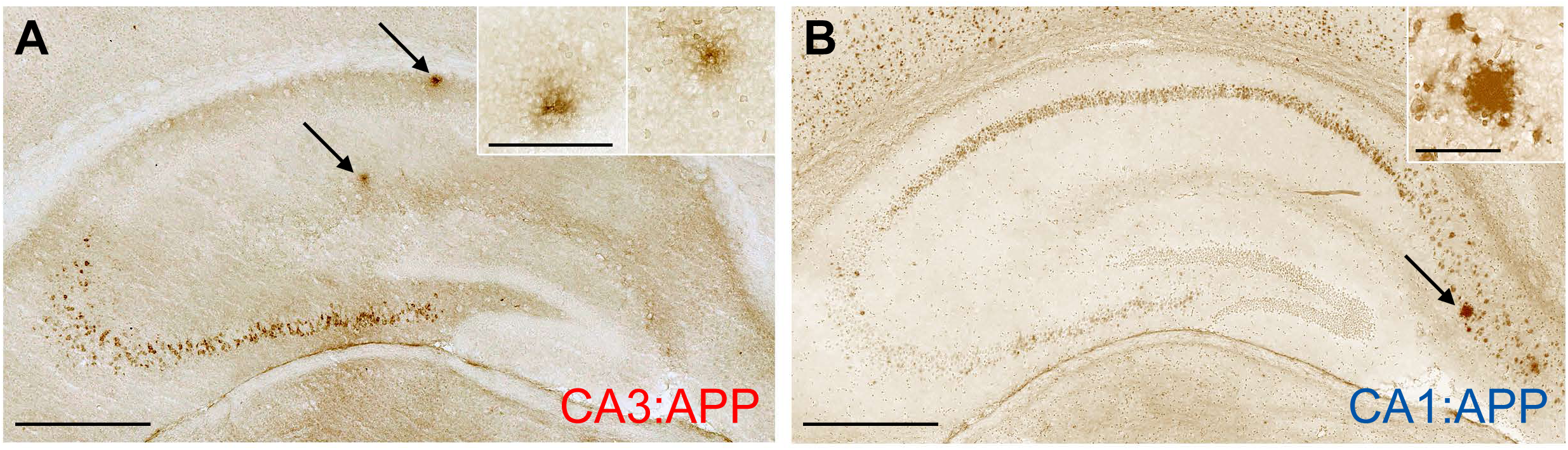
Aged APP Transgenic Mice Showed Aβ Deposits in Axonal Terminal Fields. **(a)** APP (6E10) staining of a section from a 12 month-old CA3-APP mouse. Sparse Aβ deposits were seen in both the stratum radiatum and stratum oriens (arrows), located above and below the pyramidal CA1 layer, respectively; both regions contain the synapses formed by Schaffer collaterals from CA3 neurons projecting onto the dendrites of CA1 neurons. Insets show higher magnification images of deposits. Scale bar for larger pictures: 0.5 mm, for insets: 0.1 mm. **(b)** Representative photomicrograph of a coronal section from the hippocampus of a 6 month-old CA1-APP mouse, immunostained by APP antibody (6E10). Neurons in CA1 and in subiculum are immunopositive for APP. Aβ deposits can be seen in the subiculum (arrow and inset). Note absence of Aβ deposits in the cell layer or dendritic regions of CA1 neurons (see Table 1 for numbers of analyzed mice). Scale bar: 0.5 mm.

**Table 1.**
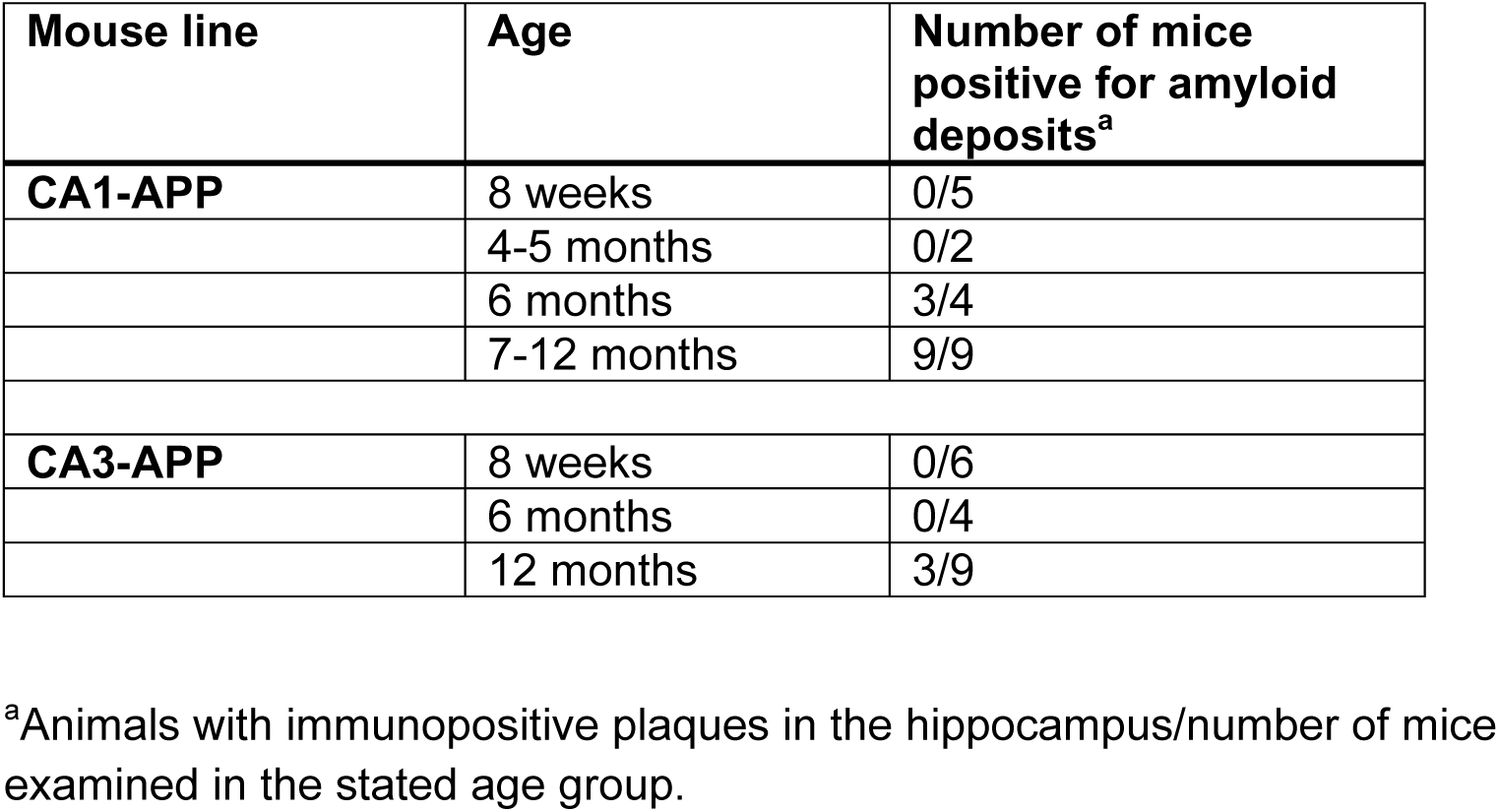
Amyloid plaque counts in CA1- and CA3-APP mice.

| **Mouse line** | **Age** | **Number of mice positive for amyloid deposits<sup>a</sup>** |
| --- | --- | --- |
| **CA1-APP** | 8 weeks | 0/5 |
|  | 4-5 months | 0/2 |
|  | 6 months | 3/4 |
|  | 7-12 months | 9/9 |
| **CA3-APP** | 8 weeks | 0/6 |
|  | 6 months | 0/4 |
|  | 12 months | 3/9 |

<sup>a</sup>Animals with immunopositive plaques in the hippocampus/number of mice examined in the stated age group.

### Synaptic Deficits Were Reversible and Due to A**β** in CA3-APP Mice

An important feature of the tetracycline system used in our CA3- and CA1-APP mice is the regulation of transgene expression. To confirm the temporal control of APP expression, doxycycline (Dox) was given to 6 week-old animals with the drug premixed into rodent chow. After two weeks of Dox treatment, there was a virtual absence of APP immunostaining in CA3-APP mice as detected by two different APP antibodies (Fig. 7A-F), indicating significant suppression of expression as predicted [21]. Western blotting of hippocampal lysates showed that Dox administration reduced APP protein levels down to basal levels present in TRE-APP mice (data not shown) [21].

**Fig. 7.**
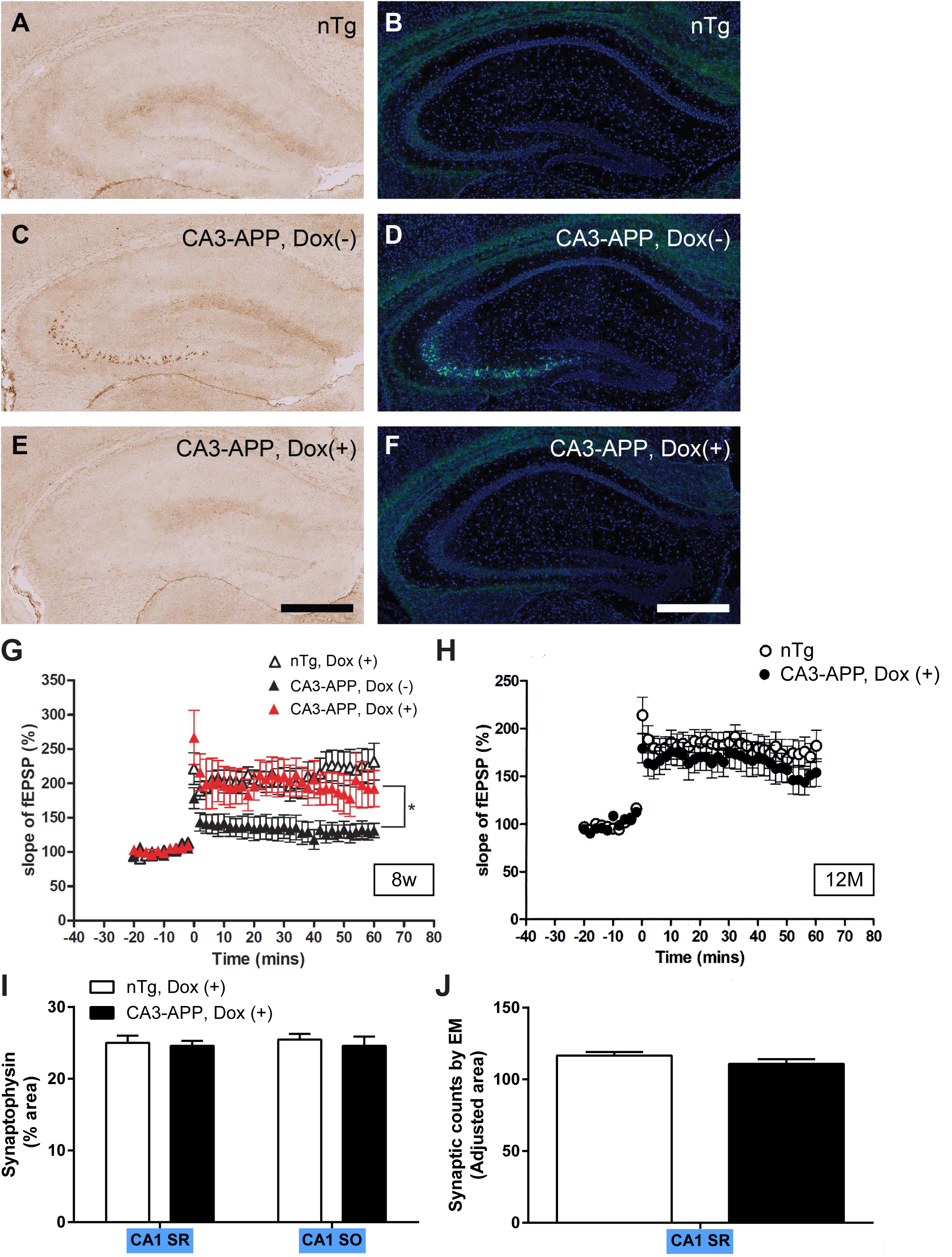
Doxycycline Treatment Suppressed APP Expression, Reversed LTP Deficits, and Prevented Synaptic Loss in CA3-APP Mice. **(a-f)** Representative photomicrographs of hippocampus immunostained with APP antibody 6E10 (a, c, e; APP: brown) or with APP antibody Y188 (b, d, f; APP: green, DAPI: blue) from (a-b) 8 week-old nTg littermates, (c-d) untreated CA3-APP mice, and (e-f) doxycycline-treated CA3-APP mice. Following 2 weeks of doxycycline treatment, APP immunostaining could no longer be detected. Scale bar: 0.5 mm. **(g)** Recordings of SC-CA1 LTP in 8 week-old mice: doxycycline-treated nTg littermates (open triangles), untreated CA3-APP mice (black triangles), and doxycycline-treated CA3-APP mice (red triangles). Doxycycline treatment reversed SC-CA1 LTP impairment in CA3-APP mice. (Values are mean ± SEM. ANOVA and *post hoc* Tukey’s test, *p<0.05, untreated vs. doxycycline-treated CA3-APP mice; nTg Dox (+): n=10 slices from 5 mice; CA3-APP Dox (-): n=13 from 6 mice; CA3-APP Dox (+): n=16 slices from 7 mice.) **(h)** 12 month-old CA3-APP mice treated with doxycline for two weeks (black circles) showed normal SC-CA1 LTP as compared to control nTg littermate controls (open circles). (Values are mean ± SEM, n=7 mice/group.) **(i)** Quantification of synaptophysin immunoreactivity in CA1 areas of 12 month-old doxycycline-treated CA3-APP mice showed no reduction compared to nTg controls. (Values are mean ± SEM, n=7 mice/group.) **(j)** Synapse quantification of synaptic counts in SBEM images from 12 month-old doxycycline-treated CA3-APP mice confirmed these results. n=2 mice/group.

Having determined that only presynaptic expression of APP resulted in reduced synaptic plasticity and increased synaptic loss, we next sought to determine whether these defects were due to APP overexpression or to Aβ. Given the similarities in results between CA1- and CA3-APP mice, we focused solely on the CA3-APP mice for these experiments. Accordingly, SC-CA1 LTP was assessed in 8 week-old CA3-APP mice treated with Dox for two weeks to suppress APP expression [21]. A restoration of the SC-CA1 LTP deficits seen previously was observed (Fig. 7G). Further, 12 month-old CA3-APP mice treated with 2 weeks of Dox exhibited equivalent SC-CA1 LTP measurements as non-transgenic controls (Fig. 7H), and transgene silencing also rescued synaptic loss in the CA1 SR and SO regions (Fig. 7I). Finally, the CA1 SR synaptic counts assessed by immunostaining of synaptic markers were confirmed using SBEM images (Fig. 7J). In all, these results confirmed the notion that the synaptic impairments were specific to the consequences of presynaptic expression of APP and also reversible.

To test whether the LTP deficits were specifically due to Aβ rather than APP or fragments generated following γ-secretase cleavage, CA3-APP mice were treated with LY-411,575, a γ– secretase inhibitor (GSI) to inhibit Aβ production [13, 73]. In 7 week-old mice, subacute treatment over 4 days with GSI led to marked elevation of APP C-terminal fragments (CTFs) in hippocampal lysates of treated mice as expected (Supplementary Fig. 5A). This elevation of CTFs was due to inhibition of γ-secretase activity, hence an accumulation of the uncleaved APP γ-secretase substrates. GSI treatment reduced hippocampal soluble Aβ40 levels by ∼75% as compared to vehicle-treated CA3-APP mice (Supplementary Fig. 5B). Importantly, GSI-treated CA3-APP and vehicle-treated nontransgenic mice showed similar SC-CA1 LTP levels (Fig. 8A), indicating that GSI treatment to inhibit Aβ production improved the LTP impairments at CA3 to CA1 synapses in CA3-APP mice. It should be noted that GSI treatment by itself depressed LTP to some degree, as drug-treated nontransgenic mice showed decreased LTP relative to vehicle-treated counterparts. In spite of this nonspecific negative effect on LTP induction, GSI was still able to increase LTP in drug-treated CA3-APP mice relative to vehicle-treated CA3-APP mice (Fig. 8B), presumably by inhibiting Aβ production. GSI treatment in 12 month-old CA3-APP mice was also able to rescue the previously observed decrease in synaptophysin immunoreactivity in the CA1 SR back to control levels (Fig. 8C). In CA1 SO, vehicle-treated CA3-APP mice in this cohort did not differ from controls in synaptophysin immunoreactivity, thereby precluding a proper assessment of GSI rescue in this region. Nonetheless, GSI treatment restored SC-CA1 LTP deficits and synapse loss, strongly suggesting that these phenotypes were directly mediated by Aβ, or, more conservatively, another APP-derived fragment derived from the myriad of APP proteolytic cleavages mediated by γ-secretase processing.

**Fig. 8.**
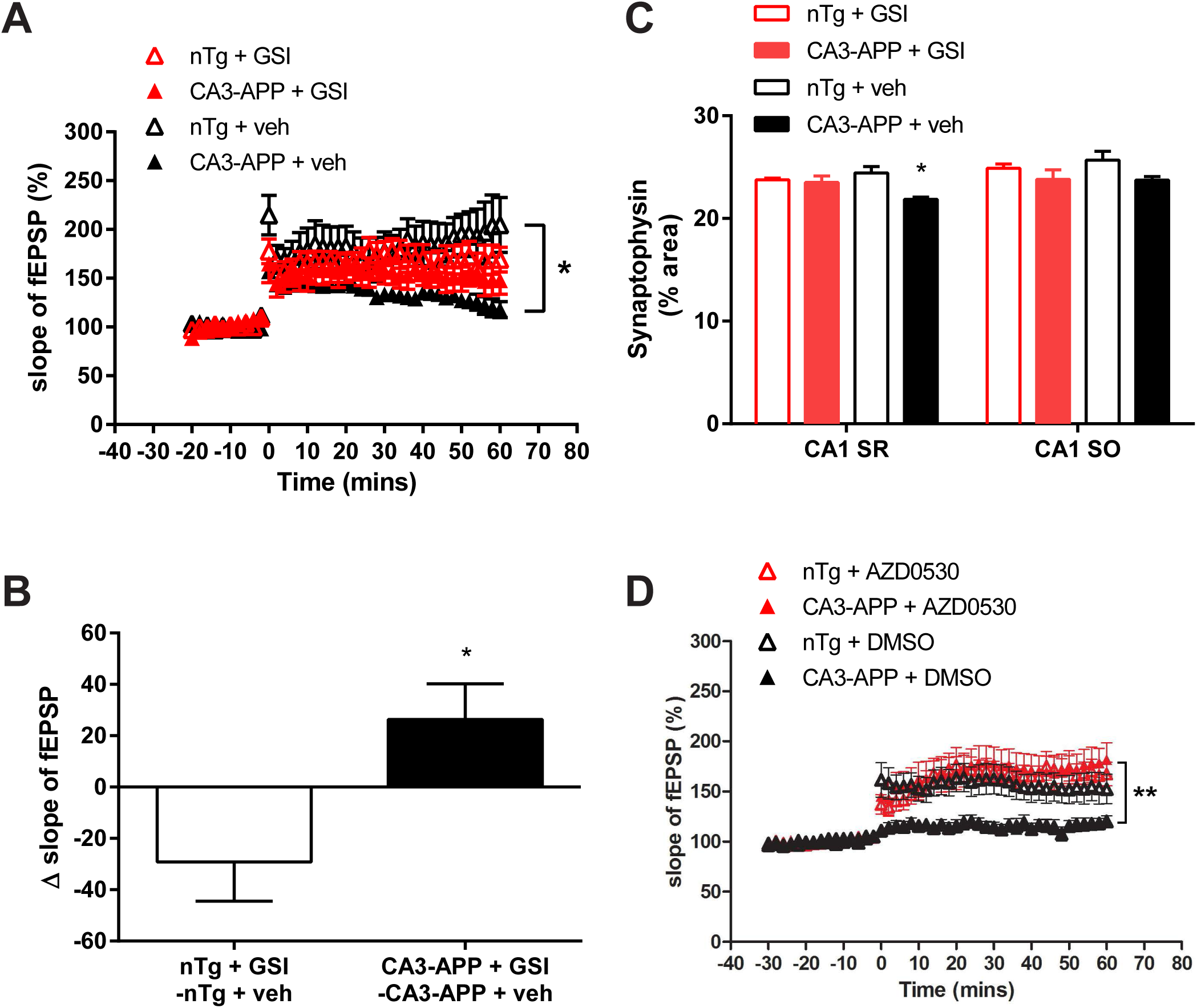
γ-Secretase and Fyn Kinase Inhibitor Reversed SC-CA1 LTP Deficits in CA3-APP Mice. **(a)** After GSI treatment, 6-8 week-old CA3-APP mice (red filled triangles) and control nTg littermates (red open triangles) showed essentially identical SC-CA1 LTP (ANOVA and *post hoc* Tukey’s test, p>0.5; nTg: n=13 slices from 6 mice; CA3-APP: n=12 slices from 6 mice), while SC-CA1 LTP was significantly reduced in vehicle-treated CA3-APP compared to vehicle-treated controls (*p<0.05; CA3-APP + veh: n=12 slices from 6 mice; nTg + veh: n=11 slices from 6 mice. Values are mean ± SEM.) **(b)** GSI treatment in nTg mice decreased LTP levels compared to nTg mice treated with vehicle alone, but increased LTP levels of CA3-APP mice vs. their vehicle-treated counterparts. Due to a confounding effect of GSI treatment which resulted in reduced LTP levels in nTg mice, mean fEPSP slopes from vehicle-treated mice were subtracted from the corresponding fEPSP slopes of GSI-treated mice. This indicates that while GSI depressed LTP relative to vehicle treatment in nTg mice, GSI increased LTP in CA3-APP mice (*p<0.01; values are mean ± SEM). **(c)** Synaptophysin immunoreactivity of 12 month-old CA3-APP and nTg mice with or without 4 days of GSI treatment. Untreated CA3-APP mice showed a significant reduction in synaptophysin immunoreactivity in CA1 SR and a trend towards a decrease in CA1 SO, as previously shown (see Fig 4B). Treatment with GSI reversed the synaptic loss in CA1 SR. (All values are mean ± SEM; n=4-6 pairs of mice/group. 2-way ANOVA with Dunnett’s multiple comparisons test, *p<0.05.) **(d)** Recordings of SC-CA1 LTP in 6 month-old CA3-APP mice (filled triangles) and nTg control littermates (open triangles) with either AZD0530 (in red) or vehicle (DMSO) treatment (in black). AZD0530 treatment given during the 30 minute basal recording period reversed LTP impairment in CA3-APP mice following stimulation of Schaffer collaterals and recording in the stratum radiatum of CA1 neurons as compared to vehicle-treated mice (**p<0.01 vehicle-treated vs. AZD0530-treated CA3-APP mice; 4 mice for each condition; nTg: 9 slices for DMSO control and 8 slices for drug treatment, CA3-APP: 8 slices for DMSO control and 10 slices for drug treatment. Values are mean ± SEM.)

As an additional test of Aβ induced toxicity, we tested the potential contribution of Src kinase Fyn to the synaptic toxicity seen in these mice because genetic and pharmacologic inhibition of Fyn have been demonstrated to protect neurons from Aβ toxicity [26, 38, 42, 60, 61]. Accordingly, we performed field recordings in acute slices of 6 month-old CA3-APP mice as before, but during the 30 minute recording period to establish baseline EPSP prior to tetanic stimulation, a Fyn kinase inhibitor, AZD0530, was added into the artificial cerebrospinal fluid (aCSF) bath [15, 42]. Intriguingly, this brief treatment was able to reverse the reduction in LTP noted previously in these mice (Fig. 8D). Thus, in the current animal model, perturbations in synaptic plasticity secondary to Aβ-induced toxicity appear to be consistent with a mechanistic role played by the Fyn kinase pathway. In summary, the LTP deficits detected in neurons situated postsynaptically to the site of APP expression appear to be mediated by APP/Aβ, while the fact that this effect can be reversed with two different Aβ-targeting treatments, GSI and Fyn kinase, suggest a primary causative role for Aβ.

## DISCUSSION

In this study, we asked whether targeting APP expression to two different groups of hippocampal neurons can provide insights into the vulnerability of synapses to Aβ-induced injury. Our results showed that APP expression can be successfully targeted preferentially to CA3 or CA1 neurons and thus provided us the opportunity to determine the pattern of synaptic dysfunction with respect to pre- or postsynaptic origin of APP expression and Aβ release. In this context, impairments in LTP and synaptic loss, the latter as assessed by both light and electron microscopy, were detected only when APP was expressed in presynaptic neurons and could be suppressed with doxycycline, supporting not only the causal role of APP but also highlighting the reversibility of synaptic defects in these mice. Importantly, LTP deficits and synapse loss were reversed with a γ-secretase or Fyn kinase inhibitor, indicating that these alterations were likely caused by Aβ. Further, because Aβ deposits were only seen in distal axonal terminal fields, we concluded that amyloid deposits likely originate primarily from the presynaptic release of Aβ into the parenchyma. Finally, the similarities in findings present in both CA1- and CA3-targeted mouse lines indicate that a presynaptic origin for Aβ-induced synaptic injury is likely a generalized phenomenon.

Our observations are compatible with the study by Harris and colleagues that selectively targeted APP to entorhinal neurons [12], although this study examined only efferent projections from the entorhinal cortex, and also with the recent report demonstrating an increase in Aβ deposits in postsynaptic sites after optogenetic stimulation to enhance neuronal activity in entorhinal cortex [75]. However, it should be noted that because the Aβ deposits in our mice, especially the CA3-APP line, were both generally sparse and occurred much later than the LTP abnormalities, these two events are unlikely to be pathophysiologically linked. This conclusion is consistent with the lack of correlation between Aβ deposits and cognition in mice or humans with AD as well as the evidence of synaptic dysfunction in APP transgenic mice many months before the onset of Aβ deposition in brain [38].

We observed that plaques only emerged at the axonal projection sites of APP-expressing neurons but not around the cell bodies of APP-expressing cells, strongly suggesting that Aβ is released axonally. There is also support for dendritic release of Aβ contributing to synaptic dysfunction in cultured neurons, as reported by other laboratories [6, 18, 23, 25, 48, 66].

Importantly, in agreement with our study, many of these experiments also showed that synaptic dysfunction does not depend on APP expression in the postsynaptic cell, as non-cell autonomous decreases in synaptic plasticity were observed when APP expression was limited to neighboring cells or to the presynaptic neuron [18, 23, 25, 66]. We speculate that the differences between our *in vivo* observations, which strongly suggest that synaptic deficits are mediated by axonal release of Aβ, and these *in vitro* studies are the high expression levels mediated by either viral delivery in organotypic slice cultures [23, 25, 66] or by overexpression of transfected APP in neuronal cultures [6, 18, 48]. Higher expression levels may enable both axonal and dendritic release of Aβ, and mask the preferential effect of synaptotoxic Aβ derived from the axonal compartment.

On the other hand, lower levels of expression *in vivo,* as used in this study, may lead to preferential release of Aβ from axonal terminals and not from dendrites, similar to what might happen physiologically in brain, consistent with our observation that plaques only emerged at the site of the synaptic projection of APP-expressing neurons but not around the cell bodies of APP-expressing cells.

Interestingly, both transgenic lines demonstrated a robust LTP deficit at relatively young ages. For instance, CA3-APP mice at 6-8 weeks of age already showed a similar reduction in LTP (by about 50%) as 4-6 month-old CA3-APP mice (Supplementary Fig. 1A & Fig. 2B) even while the number of CA3 neurons immunopositive for APP transgene expression increased from 20% at 6-8 weeks to almost 60% in the older CA3-APP mice (Fig. 1I). This suggested that there was both a possible threshold effect as well as a non-cell autonomous mechanism whereby the diffusion of Aβ peptides following release from presynaptic sites altered the function of neighboring synapses, including those that received inputs from neurons that do not express APP. Although there was a small percentage (≤10%) of CA3 neurons immunopositive for APP in CA1-APP mice, the expression remained low through 6 weeks to 6 months of age. Therefore, that small amount was likely below the threshold to cause any synaptic toxicity at CA3 to CA1 synapses, as no impairment in LTP was observed in CA1-APP mice following Schaffer collateral stimulation and recording from the CA1 stratum radiatum (Fig. 3G).

We found a pattern of postsynaptic vulnerability to APP expression when we quantified synapses either by immunohistochemistry or SBEM. There appeared to be an association between age and synaptic loss, as decreases were not detectable by immunohistochemistry in 6 month-old CA3-APP mice, but were observed in 12 month-olds (Fig. 4B). Whereas LTP deficits were detectable as early as 6-8 weeks of age, basal synaptic transmission in CA3-APP mice was unchanged at 6 months of age, but was decreased in 12 month-old CA3-APP old mice (Fig. 2C, D). In line with these observations, basal synaptic transmission deficits are commonly thought to point to a loss of functional synapses in the specific area being tested [17].

When synapse loss emerged at 12 months of age, it was nonetheless confined to subregions that receive inputs from presynaptic neurons expressing the APP transgene, such as CA1-SR and SO. In the CA3-SR layer, the reduction in synapse numbers was quite small but we interpret this as being due to the recurrent nature of these collaterals, which implies that APP can be expressed both presynaptically and postsynaptically to these axonal endings. This makes the results from analyzing recurrent collaterals less specific for the source of the toxicity than the measurements in CA1. Lastly, the absence of synapse loss in hippocampal subregions not receiving inputs from neurons expressing the APP transgene in CA3-APP mice, such as in CA1-SLM and CA3-SL, provided further confirmation that the observed reduction in synapse numbers is related to synaptic toxicity resulting from APP expression in presynaptic neurons.

Although functional abnormalities have been shown to precede outright synapse loss [12], most studies correlating the two have used surrogate markers such as synaptophysin and PSD-95, and few have taken advantage of electron microcopy to study synapse ultrastructure. Our SBEM 3D reconstruction data pointed to a reduction in spine density in 6 month-old CA3-APP mice at a time when there were no changes by immunostaining nor deficits in basal synaptic transmission (Fig. 5F). Unexpectedly, the reduction was accompanied by an enlargement of spine volumes and longer PSDs compared to non-transgenic littermate controls (Fig. 5G-I). In this regard, previous reports have found an inverse correlation between the size of synapses and synaptic density: a decrease in the density of synapses was accompanied by an increase in the size of remaining synapses [8, 49, 62]. The implications of such findings point to a possible compensatory change that takes place to increase the total synaptic contact area (TSCA) in an attempt to preserve synaptic function [50]. This compensatory mechanism may explain why basal synaptic transmission was normal in 6 month-old CA3-APP mice even though there were fewer, though enlarged, dendritic spines [62]. However, when animals reached 12 months of age, synaptic loss was more abundant and detectable by immunohistochemistry, and this may also have contributed to the now detectable abnormalities in basal synaptic transmission observed in these older mice.

The reversal of SC-CA1 LTP impairment in CA3-APP mice after short-term treatments of doxycycline (Fig. 7G) or especially GSI (Fig. 8A) in 2 month-old mice implied that Aβ was the likely toxic stimulus released by presynaptic neurons and that the deficits are reversible at this age. Synapse loss at the CA1 SR or SO was also reversed by doxycycline and GSI treatment in 12 month-old CA3-APP mice (Fig. 7I-J, Fig. 8C), suggesting that even structural deficits remained reversible, which perhaps was not predicted at the outset. In addition to Aβ, APP generates multiple proteolytic fragments, such as secreted derivatives (sAPPα, sAPPβ, and sAPPη) or cell retained fragments (CTFα, CTFβ, and CTFη) [40, 70], that have been shown to play pleiotropic roles, primarily in the cultured setting. It is difficult to definitively exclude the possibility that other non-Aβ APP-derived fragments can also induce synaptic toxicity. Notably however, SC-CA1 LTP was also restored to normal levels by a brief incubation with a Fyn kinase inhibitor in CA3-APP mice (Fig. 8D), consistent with the proposed role of Fyn kinase in mediating Aβ-induced toxicity [4, 31, 45]. In view of the findings from the LTP experiments after treatments with doxycycline, GSI, and Fyn kinase inhibitor, we conclude that the most parsimonious explanation is that the synaptic toxicity observed in CA3- and CA1-APP mice is derived primarily from the release of Aβ from presynaptic terminals.

Selective regional loss of neurons is a consistent feature in virtually all neurodegenerative and many neurologic disorders, such that the pattern of cell damage can be grouped by, for example, anatomic regions or neurotransmitter systems. Yet, satisfactory explanations for these characteristic patterns of neuronal injury have been elusive. The pattern of synaptic dysfunction described in this study indicates that presynaptic Aβ release altering postsynaptic mechanisms plays a critical role in Aβ-mediated synaptic injury *in vivo*. In this way, it is consistent with the spread of pathology that is increasingly being described in various neurodegenerative diseases, especially in tauopathies and synucleinopathies [16, 22, 35, 47, 74]. In conclusion, the approach taken here may offer the opportunity to isolate distinct neuronal populations to interrogate in greater depth the vexing problem of selective neuronal vulnerability in brain.

## MATERIALS AND METHODS

### Animals

All animal procedures were approved by the Institutional Animal Care and Use Committee of the University of California San Diego. tetAPP mice (line 102) [21] were kindly provided by Dr. Joanna Jankowsky. α-CaMKII promoter Cre mice (line “B6.Cg-Tg (CaMKIIa-cre) T29-1Stl/J”) [58] and Grik4 promoter Cre mice (line “C57BL/6-Tg (Grik4-cre) G32-4Stl/J”) [39] were generously provided by Dr. Susumu Tonegawa. ZtTA mice are engineered such that the tTA expression cassette inserted in the Rosa26 locus is flanked by two loxP sites [33]. All lines except ZtTA were congenic in C57BL6 background. ZtTA mice originated in mixed C57BL/6, FVB/N, and CD-1 backgrounds and were maintained in a mixed background [33]. To obtain the desired genotype, mice homozygous for the ZtTA transgene were first bred with TRE-APP heterozygous transgenic mice. The resulting ZtTA/TRE-APP bigenic mice were crossed with site-specific Cre (CA1 or CA3) mice. Male triple transgenic breeders were bred to CD-1 female mice to obtain triple transgenic offspring in mixed strain backgrounds. For controls, littermates lacking one or more transgenes were used except those positive for TRE-APP transgene were discarded to avoid any basal APP expression from the minimal tetracycline responsive promoter that might confound the analyses. The number of animals analyzed in the reported experiments are summarized in Table 1 and Supplementary Table 1. Mice were deeply anesthetized with isoflurane and perfused transcardially with ice-cold phosphate-buffered saline (PBS) before the brains were removed and frozen for subsequent analyses; for immunofluorescent staining, brains were removed and fixed in 4% paraformaldehyde (PFA) for 24 hours.

### Drug treatments

Doxycycline (Dox) was added into rodent chow through custom formulation at a concentration of 50-100 ppm antibiotic (TestDiet, Richmond, IN, US) [21].

To inhibit γ-secretase activity, LY-411,575 (kind gift from Dr. Todd Golde, University of Florida), was formulated as a 5 mg/mL solution in 50% polyethylene glycol, 30% propylene glycol, 10% ethanol and diluted 1:10 with 20% hydroxyl-propyl-β-cyclodextrin immediately before use [30]. Mice were injected subcutaneously once daily for 5 days at a dose of 5 mg/kg or with vehicle and sacrificed 5 hours after the last injection.

Fyn inhibitor, AZD0530, was dissolved in DMSO and diluted 1:2000 with ACSF just before the experiment. 2 μ of AZD0530 was perfused for 30 min prior to LTP induction [15, 41].

### Acute Hippocampal Slice Electrophysiology

Extracellular field potential recordings were assessed in three groups of animals: from between 6 to 8 weeks of age, from 4 to 6 months of age and in 12 month-old mice. A minimum of 10 slices were measured from 5-7 mice for each cohort of animals. A maximum of two mice, one experimental and one littermate control, blinded to the tester, were sacrificed and analyzed each day. Acute hippocampal slices were prepared following euthanasia under isoflurane anesthesia, according to the protocol described previously [59]. In brief, the brain slices were kept at room temperature for 1-2 hours before transfer to a recording chamber in artificial CSF (ACSF) containing 125 mM NaCl, 2.4 mM KCl, 1.2 mM NaH_2_PO_4_, 1 mM CaCl_2_, 2 mM MgCl_2_, 25 mM NaHCO_3_, and 25 mM glucose. In the recording chamber, the slices were perfused with oxygenated ASCF containing 2 mM CaCl_2_ and 1 mM MgCl_2_. Extracellular recordings of excitatory postsynaptic potentials (fEPSPs) were obtained from three hippocampal regions (see

below) using microelectrodes filled with extracellular recording solution. A concentric bipolar stimulating electrode was placed in the dendritic fields receiving the axonal projections being stimulated. Basal synaptic transmission was assessed by comparing the input and output relationship of the recorded field potentials. For each animal, we recorded the fiber volley amplitude and initial slope of the fEPSP responses for a range of stimulation intensities from 100 to 900 µA. The strength of synaptic transmission was quantified by measuring the peak amplitude of the fiber volley (input) and the initial slope of the fEPSP (output). The input output (I/O) curve was obtained by plotting all individual fEPSP slope and fiber volley values, and the data was then fitted by linear regression analysis. For examining synaptic plasticity, the stimulus intensity was adjusted individually for each experiment to produce fEPSPs which are 30-40% of the maximal responses that could be evoked. Short-term and long-term synaptic modifications were measured by paired-pulse facilitation (PPF) and long-term potentiation (LTP), respectively. PPF were evoked with interstimulus intervals of 50 ms at submaximal stimulus intensities. PPF is expressed as the ratio of the peak amplitude of fEPSP (second stimulation)/fEPSP (first stimulation). LTP were induced by four tetani delivered 20 sec apart, each at 100 Hz for 1 sec after a 20 minute baseline period. fEPSPs were monitored for 60 minutes after tetanus. Synaptic plasticity was assessed at three sites: (1) stimulation of Schaffer collateral/commissural pathway and recorded from CA1 stratum radiatum, (2) stimulation of mossy fiber from granule cells of the dentate gyrus and recorded from the stratum lucidum of CA3 neurons, and (3) stimulation of CA1 axonal projections and recorded from the substratum radiatum in subiculum. In the last condition, hippocampal slices were obtained from horizontal rather than coronal sections.

Because of the complex circuitry of the CA3 area, the metabotropic Glutamate receptor (mGluR) II agonist 2-(2,3-dicarboxy-cycloproply)glycine (DCG-IV) was used at the end of experiments to verify that LTP was induced by mossy fiber inputs (data not shown) [24]. The raw data was collected and analyzed by using pClamp 10 software (Molecular Devices, Sunnyvale, CA, USA).

### Immunohistochemistry

Immunostaining for APP/Aβ was carried out on 10 µ m brain tissue cryosections post-fixed with 4% PFA. For plaque visualization, sections were treated with 88% formic acid prior to staining. Biotinylated 6E10 mouse monoclonal antibody (Signet, Dedham, MA) and Y188 rabbit monoclonal antibody (Epitomics, Burlingame, CA) were used for immunolocalization of APP. 6E10 antibody is human-specific and recognizes the N-terminus of Aβ within APP, thus detecting both APP and Aβ deposits. A polyclonal rabbit antibody (69D) generated against Aβ peptide which failed to recognize APP on immunohistochemistry was used to immunostain Aβ deposits. Appropriate HRP-tagged secondary antibodies were used, and signal development was accomplished using the Vectastain Elite ABC HRP Kit and the DAB Peroxidase Substrate Kit (VectorLabs); Nuclear Fast Red (VectorLabs) was used for counterstaining if desired. For each case, 5–15 sections taken from anterior to posterior hippocampus were assessed by immunohistochemistry. For synaptic marker staining, a vibratome was used to cut 30 µ m sections from brains fixed in 4% PFA. Primary antibodies used were: biotinylated mouse anti-6E10 (1:1000, Covance), mouse anti-synaptophysin (1:500, Dako), rabbit anti-PSD-95 (1:500, Epitomics), and mouse anti-Pcp4 (1:1000, Sigma). For synaptophysin and PSD-95 immunostaining, primary antibodies were incubated for three nights at 4°C followed by incubation with donkey anti-mouse or anti-rabbit IgG secondary antibodies (1:1000, Life Technologies). Fluorescent images were acquired with an Olympus FV1000 at 2048 x 2048 pixel resolution using a 40x objective lens. At least three random images were taken for each of the hippocampal subregions analyzed (CA1 SR, SO, SLM; CA3 SR, SL, SO) and between 5-7 sections were imaged per mouse. The number of pixels containing synaptophysin or PSD-95 immunoreactivity was measured per unit area above a minimum threshold intensity determined in the thresholding function of Image J (National Institutes of Health): histograms of pixel brightness reflecting a bimodal distribution of background peak (mode) plus an SD above background were determined. Synaptophysin and PSD-95 data were expressed as a percentage of the sampling area occupied by the staining [36]. Results were expressed as the average of experiments ± SD (or SEM).

### Serial-block electron microscopy

SBEM data was acquired at the National Center for Microscopy and Imaging Research (NCMIR) and analyzed using IMOD software (http://bio3d.coloroda.edu/imod/). Mice were transcardially perfused with 0.15M cacodylate buffer containing 2 mM CaCl, 2% formaldehyde, and 2.5% glutaraldehyde. Brains were post-fixed overnight at 4°C and coronal slices (200 µm) were sectioned on a vibratome microtome, washed overnight in 0.15M cacodylate buffer, and then osmicated in 2% osmium tetraoxide /1.5% potassium ferrocyanide for 1 hour, 0.5% aqueous thiocarbohydrazide for 20 minutes, and 2% osmium tetraoxide for 30 minutes. Slices were extensively washed with water in between the stains. After overnight incubation in 2% uranyl acetate solution, the sections were dehydrated through graded ethanol and acetone washes.

Slices were infiltrated with Durcupan ACM epoxy resin and cured in the oven at 60°C for 48 hours. Small regions approximately 1mm x 1mm were cut from the sections with a razor blade and mounted on Aclar. A glass knife was used to remove plastic from the face of the specimen. The specimen was removed from the Aclar and mounted to an aluminum scanning electron microscope (SEM) rivet with conductive silver epoxy (Ted Pella) with the exposed tissue face in contact with silver epoxy. The specimens were imaged in either a Zeiss Merlin or Zeiss Merlin Compact SEM equipped with a Gatan 3View system. Volumes were collected with 50 nm step size with each volume consisting of 25k x 25k images for each slice where the pixel size was between 3-7 nm. The acquired volumes were stacked and aligned by cross-correlation using IMOD [28]. Neuropil spine density analysis was performed on randomly selected sub-volumes of neuropil using the stereology tool within IMOD. For synaptic counts, SBEM images were used to analyze the number of synapses blinded to genotype. A synapse was defined as an electron dense post-synaptic density area juxtaposed to presynaptic terminals filled with synaptic vesicles. Synaptic counts were adjusted by area. For dendrite reconstruction, CA1 SR dendrites and spines were manually segmented using IMOD. Dendrites were manually traced for approximately 10 µm after they branched from large diameter primary dendrites [68].

### Western blotting

Immunoblotting was carried out from lysates prepared from PBS-perfused brains and sonicated in five-fold volumes of 1% CHAPSO with protease inhibitor cocktail (Sigma-Aldrich). After centrifugation at 100,000×*g* for 1 hr, supernatants were 1:20 diluted with RIPA buffer. Protein concentrations were measured by BCA Protein Assay Reagent (Thermo Scientific). Lysate samples were fractionated by 8% Tris-Glycine (to detect full-length APP) or 12% Bis-Tris (to detect APP-CTFs) gel electrophoresis, transferred to nitrocellulose membranes and blocked in Tris-buffered saline (TBS) containing 2% Tween-20 (TBS-T) and nonfat dry milk powder. The membranes were incubated overnight with 6E10 (1:5,000), CT15 (1:10,000), a rabbit polyclonal antibody recognizing the APP C-terminus [54], or mouse monoclonal anti β-tubulin antibody E7 (1:20,000; Developmental Studies Hybridoma Bank). Immunoreactive species were visualized by SuperSignal West Pico Chemiluminescent Substrate (Thermo Scientifc) and signal intensity was quantified densitometrically with NIH ImageJ software.

### Quantification of Aβ

For quantification of Aβ40 levels in brains of transgenic mice, hippocampi were homogenized in 1% CHAPSO or 2% SDS and detergent-soluble supernatants were analyzed by Mesoscale Discovery human Aβ40 kit (MSD) as per manufacturer’s instructions.

### Statistical Analyses

Statistical analyses for all experiments were performed with GraphPad Prism 5 as indicated in the figure legends or text. Experimenters were blinded with respect to genotype for all measurements.

## COMPLIANCE WITH ETHICAL STANDARDS

Animals were maintained in an AAALAC-accredited facility in compliance with the Guide for the Care and Use of Laboratory Animals. All animal procedures were approved by the University of California San Diego, Institutional Animal Care and Use Committee.

## ACKNOWLEDGEMENTS

This work was supported in part by NIH grants NS84324 (EHK, SL), AG32179 (EHK), NS097772 (SL), NS102915 (SL), NMRC/STaR/009/2012 (EHK), the donors of the Alzheimer’s Disease Research program of the BrightFocus Foundation A2015595F (KC), “La Caixa” Foundation Fellowship (EV-O), and AG 005131 (KK, S-HT). Light microscopy was performed at the UCSD School of Medicine Microscopy Core, which is supported by an NINDS P30 grant (NS047101). Electron microscopy was performed at the NCMIR, which is supported by an NIGMS P41 grant (GM103412). We are grateful to Dr. Susumu Tonegawa and Dr. Joanna Jankowsky for the gift of transgenic mouse lines. We thank the late Dr. Stephen Heinemann for advice and valuable discussions leading to this project, Dr. Wei Guo, Kaivon Sobhani, Cheng-Lin Shaw, Erin Teramoto, and Ivy Dang for technical assistance, and Dr. Kazunari Miyamichi for critical reading of the manuscript.

