## Supplemental Legends and Figures for "Aβ-induced synaptic injury is mediated by presynaptic expression of amyloid precursor protein (APP) in hippocampal neurons"

### SUPPLEMENTARY FIGURE LEGENDS

**Fig. S1**

**Pre-synaptic APP Expression Impaired Synaptic Plasticity But Not PPF**

**(a)** 6-8 week-old CA3-APP mice (black triangles) showed significant impairment in LTP compared to control non-transgenic littermates (nTg, open triangles; ***$p<0.005$) when APP was expressed in CA3 neurons and field potentials were measured in stratum radiatum of CA1 neurons following stimulation of Schaffer collaterals (SC-CA1 LTP).

**(b)** Using the same stimulation and recording paradigm as in (A), no PPF deficits were observed in 4-6 month-old CA3-APP mice with 50, 100, 200, 300 and 400 ms intervals.

**(c-d)** At 4-6 months of age, there were no PPF deficits in CA3-APP mice [$p>0.5$, same stimulation/recording paradigm as in (A)], nor at 12 months of age (D).

**(e)** PPF deficits were not observed in 4-6 month-old CA1-APP mice when CA1 axons were stimulated and recording was performed in the subiculum ($p>0.5$).

**Fig. S2**

**Post-synaptic APP Expression Did Not Impair PPF**

**(a)** No PPF deficits were observed in CA3-APP compared to nTg mice when mossy fibers were stimulated and field potentials were measured in CA3 stratum lucidum ($p>0.5$).

**(b)** No PPF deficits were observed in CA1-APP mice following Schaffer collateral stimulation and measurement of field potentials in stratum radiatum of CA1 neurons.

**Fig. S3**

**PSD-95 immunoreactivity in CA3-APP mice is only decreased in regions receiving axonal projections from the APP-expressing CA3 neurons**

**(a)** Immunostaining of PCP4 (green), a CA2 marker, together with human APP, stained with 6E10 (red). Scale bar 50 µm.

**(b)** Percent area of PSD-95 immunoreactivity in CA1 SR, CA1 SO and CA3 SR in 6 and 12 month-old CA3-APP mice. 12 month-old CA3-APP mice had decreased immunoreactivity in all three areas as compared to control littermates. All values are mean ± SD; n=9 mice/group.
**(c)** Percent area of PSD-95 immunoreactivity in CA1 SLM and CA3 SL of 12 month-old CA3-APP mice was unchanged compared to control littermates.

**Fig. S4**

**Aβ Deposits in CA1- and CA3-APP mice**

**(a,c-f)** Representative photomicrographs of hippocampus immunostained with APP antibody 6E10 or (b) the plaque-specific Aβ antibody 69D; insets contain higher magnification images of deposits.
**(a-d)** CA1-APP mice, (c-d) slices were counterstained with Nuclear Fast Red, (d) higher magnification image of plaques located in the cortex of 10 month-old CA1-APP mice. Arrowheads indicate staining consistent with cerebral amyloid angiopathy (CAA), or vascular deposition of Aβ. Note the increase in subicular and cortical deposits in 10 month-old vs. 7 month-old CA1-APP mice.
**(e-f)** Slices from the posterior hippocampus of CA3-APP mice containing solitary deposits in the dendritic fields of CA1 neurons. Scale bar for larger pictures: 0.5 mm, for insets: 0.1 mm.

**Fig. S5**

**γ-Secretase Inhibitor Treatment Suppressed Aβ Production in CA3-APP Mice**

**(a)** Representative Western blots of full-length APP (FL) and APP C-terminal fragments (CTFs) in hippocampi of CA3-APP mice with and without GSI treatment. 6E10 and CT15 antibodies were used for detection. APP-CTFs were markedly elevated after 4 days of GSI treatment in 7 week-old mice.
**(b)** Quantification of Aβ40 levels in hippocampi by MSD assay demonstrated ~70% reduction in Aβ40 levels following GSI treatment. (***$p < 0.001$ by Tukey's posthoc test after one-way ANOVA; n=3 mice/group.)

**Table S1**

**Numbers of mice used for electrophysiology experiments and synapse quantification**

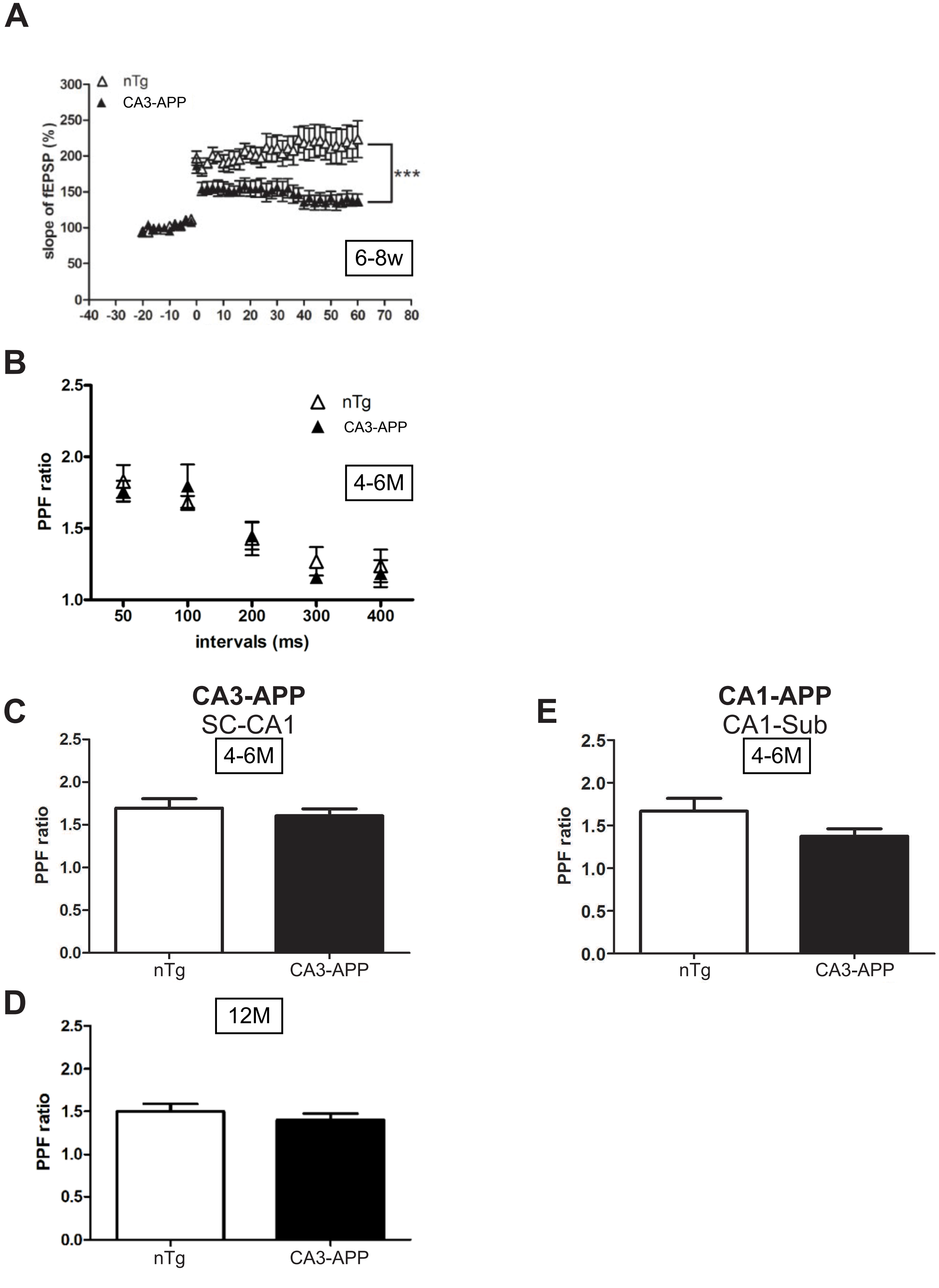


Supplementary Fig. 1

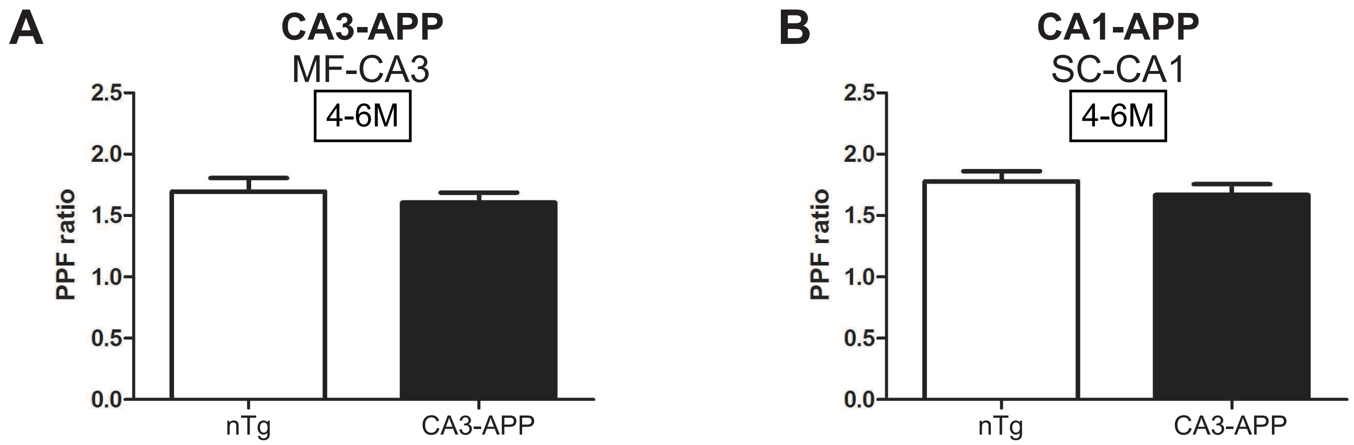


Supplementary Fig. 2

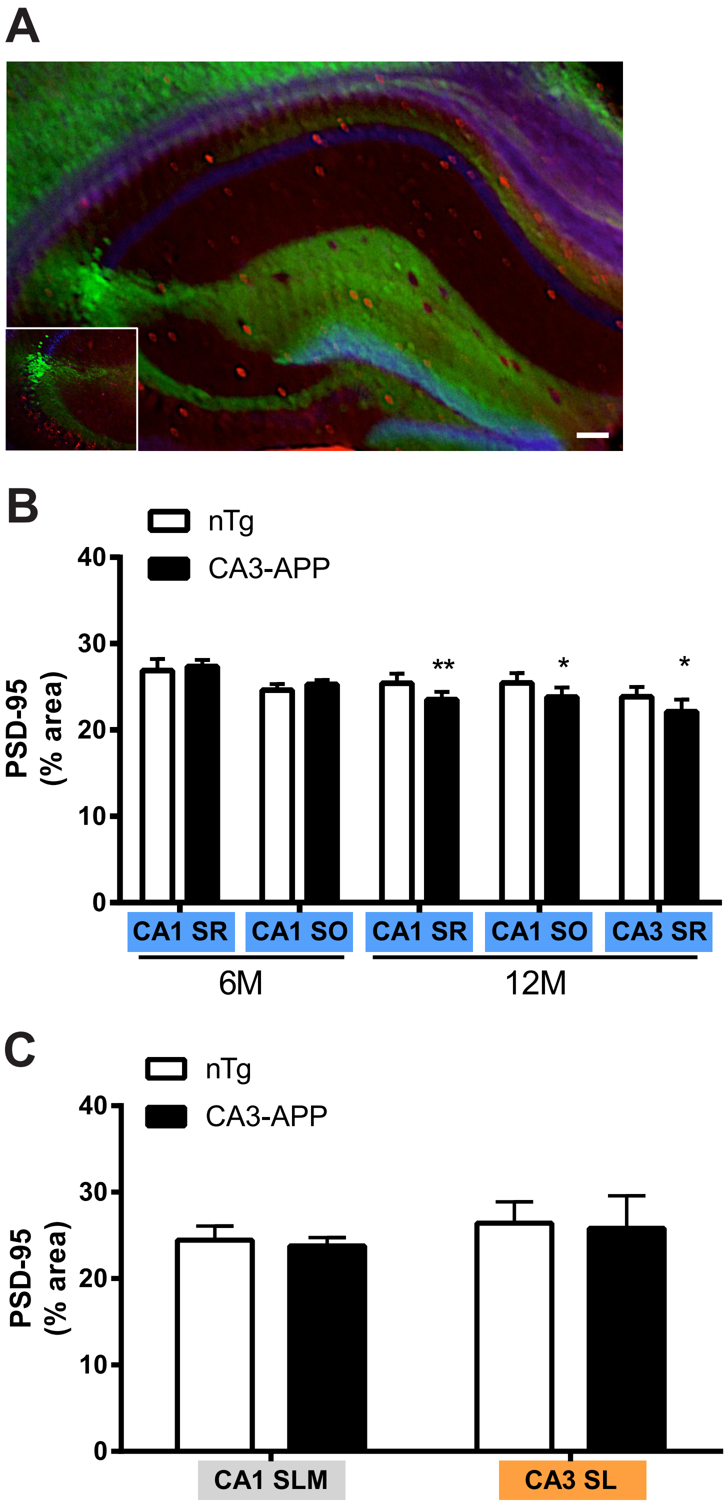


Supplementary Fig. 3

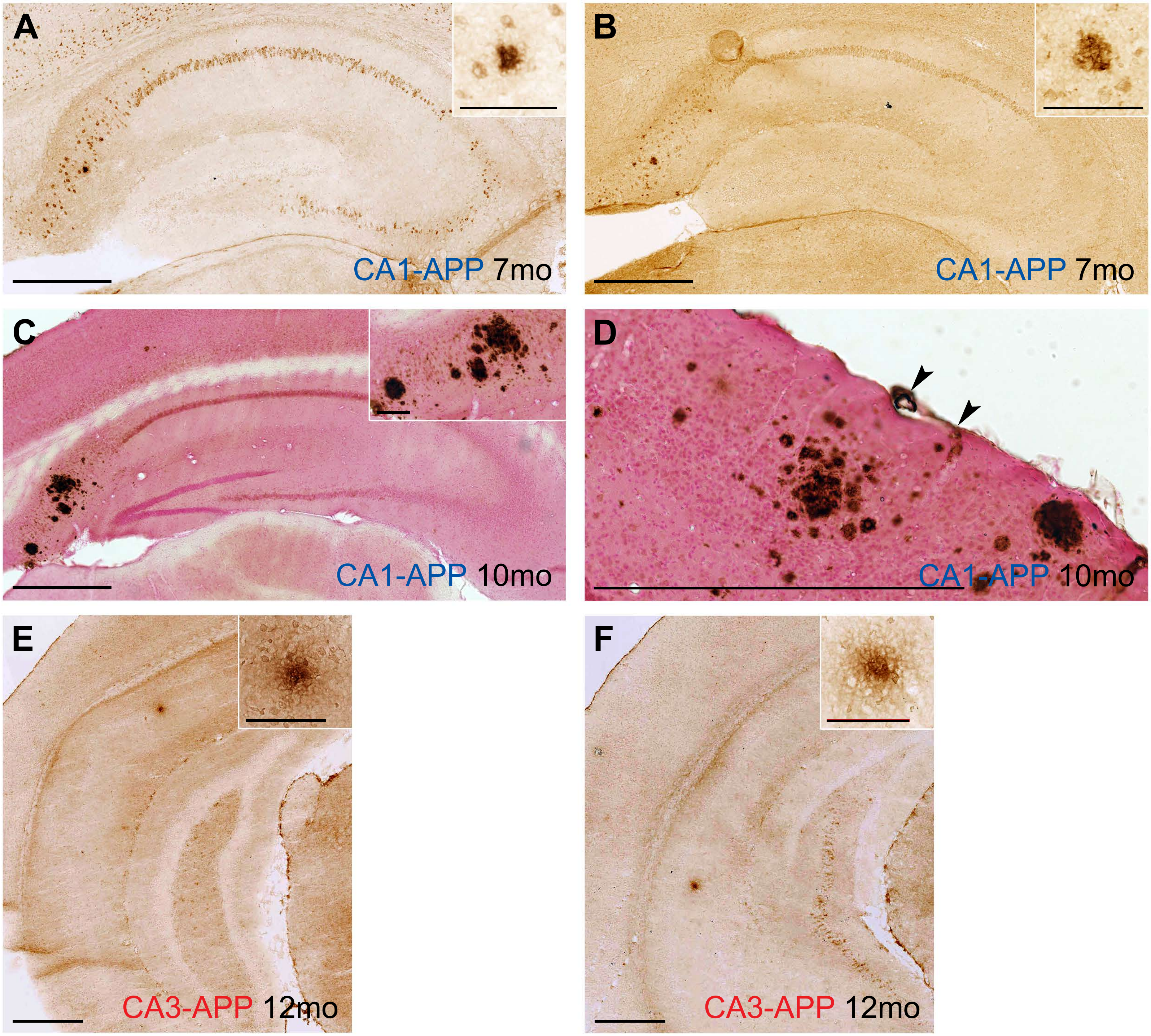


Supplementary Fig. 4

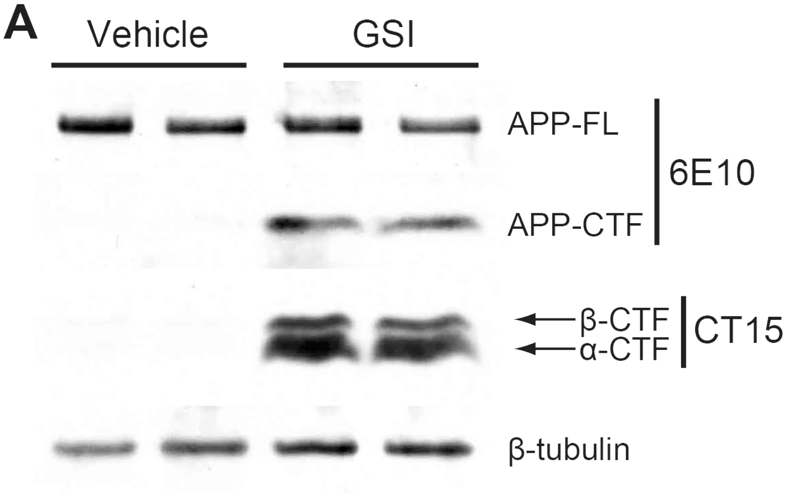


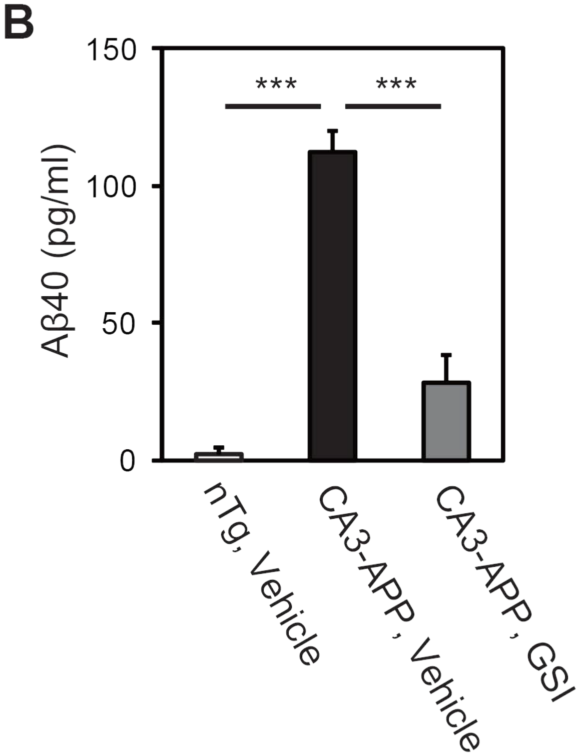


Supplementary Fig. 5

### SUPPLEMENTARY TABLE 1

| **Electrophysiology** | | | |
|---|---|---|---|
| **Mouse line** | **Age** | **Genotype** | **No. mice** |
| CA1-APP | 6-8 weeks | Transgenic<br>Control | 8<br>8 |
| | 4-6 months | Transgenic<br>Control | 12<br>13 |
| CA3-APP | 6-8 weeks | Transgenic<br>Control | 6<br>5 |
| | 4-6 months | Transgenic<br>Control | 13<br>10 |
| | 12 months | Transgenic<br>Control | 7<br>7 |
| CA3-APP + Doxycycline | 8 weeks | Transgenic<br>Control | 13<br>5 |
| CA3-APP + GSI | 8 weeks | Transgenic<br>Control | 5<br>6 |
| CA3-APP + Fyn kinase inhibitor | 6 months | Transgenic<br>Control | 8<br>8 |

| **Synapse quantification** | | | |
|---|---|---|---|
| **Mouse line** | **Age** | **Genotype** | **No. mice** |
| CA1-APP | 6 months | Transgenic<br>Control | 9<br>9 |
| CA3-APP | 6 months | Transgenic<br>Control | 9<br>9 |
| | 12 months | Transgenic<br>Control | 9<br>9 |
| CA3-APP + Doxycycline | 12 months | Transgenic<br>Control | 7<br>7 |
| CA3-APP + GSI | 12 months | Transgenic<br>Control | 5<br>7 |
| CA1-APP by SBEM | 6 months | Transgenic<br>Control | 2<br>2 |
| CA3-APP by SBEM | 6 months | Transgenic<br>Control | 2<br>2 |
| | 12 months | Transgenic<br>Control | 2<br>2 |
